# Ncbe is the main basolateral Na^+^ loading mechanism of the choroid plexus epithelium

**DOI:** 10.64898/2026.08.24.745951

**Authors:** Lasse Messell Desdorf, Sigurd Kassow Morsby, Laura Øllegaard Johnsen, Nicoline Stærke Jensen, Christian Andreas Hübner, Helle Hasager Damkier, Jeppe Praetorius

**Affiliations:** Department of Biomedicine, Aarhus University, Denmark; Department of Chemistry, Aarhus University, Denmark; Institute of Human Genetics, University Hospital Jena, Germany

## Abstract

Cerebrospinal fluid (CSF) provides a specialized extracellular environment for the central nervous system, which is predominantly produced by the choroid plexus, a highly vascularized epithelial structure whose ion transport processes are fundamental to CSF secretion, composition, and homeostasis. The mechanisms of Na^+^ entry into choroid plexus epithelial cells (CPECs) from the interstitial side remain disputed. The *slc4a10* gene product encoding the Na^+^ dependent Cl^-^/HCO_3_^-^exchanger, Ncbe, was suggested as a key transport mechanism based on its impact on the cell’s Na^+^-dependent regulation of intracellular pH and its basolateral membrane expression. The current study was undertaken to directly assess the contribution of Ncbe to the Na^+^ uptake into CPECs. Intracellular Na^+^ was recorded by fluorometry using the Na^+^ probe Sodium Binding Fluorescent Indicator in clusters of CPECs with access to both the luminal and basolateral membranes. Removal of extracellular Na^+^ reduced the apparent *ex vivo* intracellular [Na^+^] to ∼5 mM from a baseline of ∼43 mM in the absence of CO_2_/HCO_3_^-^ and ∼54 mM in the presence of CO_2_/HCO_3_^-^. Flame photometry estimated the intracellular [Na^+^] *ex vivo* to ∼28 mM. The CO_2_/HCO_3_^-^-dependent rate of [Na^+^] recovery amounted to ∼53% of the total recovery rate upon re-addition of Na^+^. Experiments with access to only the luminal membrane show a [Na^+^] recovery of a similar rate as observed in the absence of CO_2_/HCO_3_^-^ in the clusters. The CO_2_/HCO_3_^-^-independent [Na^+^] recovery was inhibited to ∼50% by the NKCC1 inhibitor bumetanide and to ∼30% by the TRPv4 inhibitor RN1734. NHE contributed to a minor extent to the CO_2_/HCO_3_^-^-independent transport. The HCO_3_^-^ transport inhibitor DIDS, however, inhibited the total [Na^+^] recovery rate to ∼50%, indicating a role for Ncbe rather than NBCn1 in the cellular [Na^+^] recovery. Indeed, docking of DIDS into Ncbe and NBCn1 indicated that both proteins can accommodate the binding of DIDS. However, the orientation of the DIDS poses in Ncbe suggests a binding mode more similar to that found in the Anion Exchangers (SLC4A1-3), which seems to accommodate the covalent-type docking more than NBCn1. The Ncbe inhibition by DIDS was supported by the rate of [Na^+^] recovery that was significantly higher in CPECs from Ncbe-wt than Ncbe-ko mice in the presence of CO_2_/HCO_3_^-^. As both NKCC1 and TRPv4 are localized to the luminal membrane, the findings collectively suggest that Ncbe is the most prominent mechanism for Na^+^ entry into CPECs expressed at the basolateral side. We suggest Ncbe as the rate-limiting mechanism in the vectorial Na^+^ transport driving CSF secretion.

## Introduction

The majority of the intra-cerebroventricular cerebrospinal fluid (CSF) is secreted by choroid plexus epithelial cells (CPECs) through a high-rate transport process driven by active transcellular Na^+^ movement (1, 2). The net secretory process is a concerted action of multiple plasma membrane transport proteins specialized for transcellular and, in some cases, paracellular movement of ions and water. Figure 1 shows the transporters considered important for CSF secretion by the choroid plexus, but for some of these proteins direct evidence for their involvement in cellular Na^+^ transport is lacking. The lack of knowledge is especially apparent for basolateral Na^+^ entry into CPECs, which is considered the rate-limiting step in secretion here (1, 2). Basolateral Na^+^ entry was previously thought to be carried by an amiloride-sensitive Na^+^/H^+^ exchange mechanism (3–5), which would work in parallel with the anion exchanger AE2 to bring net Na^+^ and Cl^-^ into the cells from the interstitial side of the epithelium. Secreted HCO_3_^-^ was formed inside the epithelium, catalyzed by cytosolic carbonic anhydrases, as judged by the sensitivity of HCO_3_^-^ secretion to acetazolamide (6, 7). Previous studies, however, demonstrated minimal NHE immunoreactivity (8, 9) and even NHE peptide detection by mass spectrometry was challenging (10). Nevertheless, NHE activity has long been described in the choroid plexus as assessed by the dimethadione method (4). In our hands, both NHE1 and NHE6 immunoreactivity, as well as NHE activity, were unexpectedly confined to the luminal membrane domain of the CPECs from mice and only demonstrated the basolateral membrane domain in CPECs from Ncbe-ko mice (9). Thus, the main candidate for mediating Na^+^ entry from the interstitial side to sustain CSF secretion was no longer supported at the molecular level.

**Figure 1.**
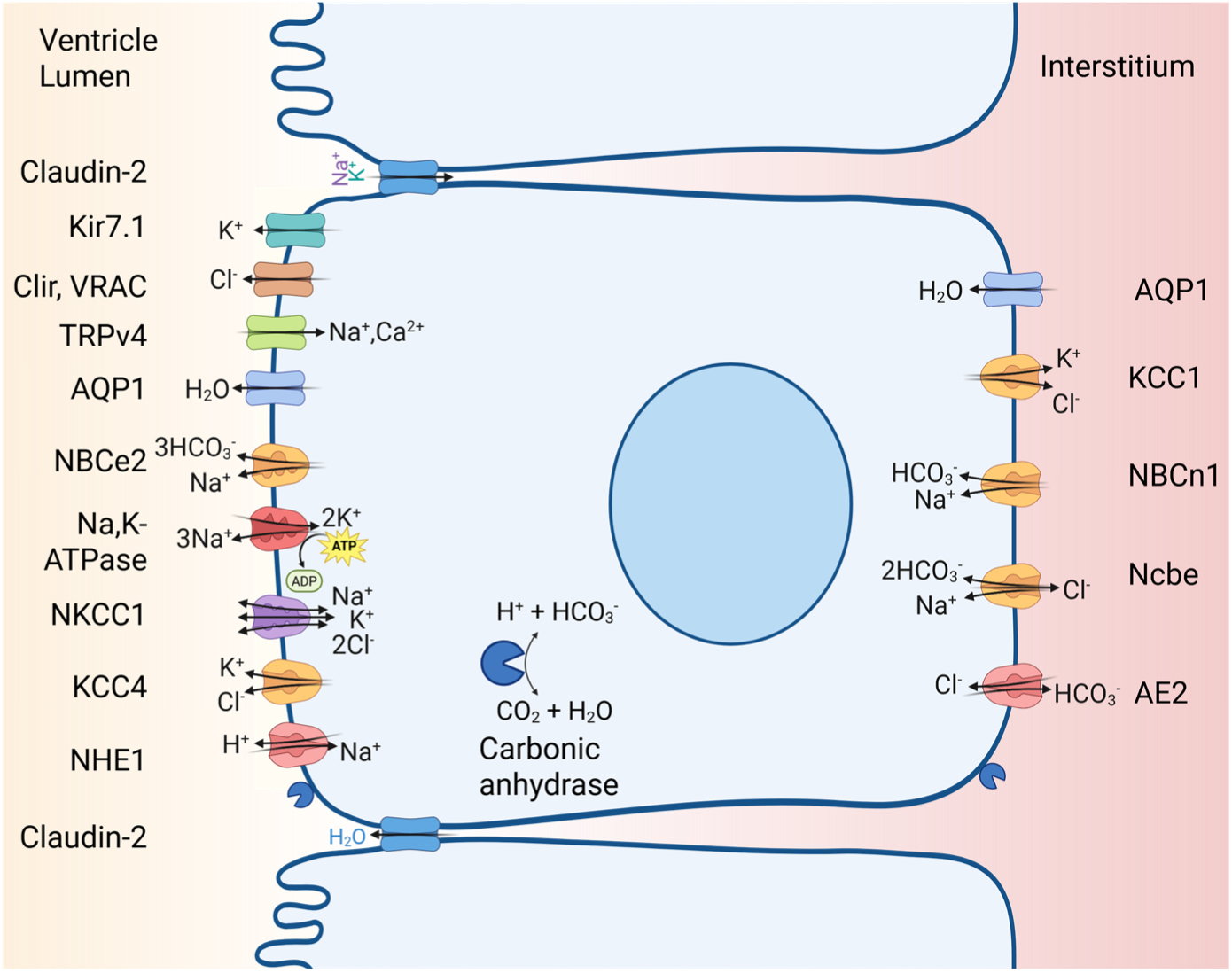
Schematic representation of the main transport pathways across a choroid plexus epithelial cell. The cerebroventricular lumen is shown on the left, and the interstitium facing the blood vessels is on the right. The individual types of transport proteins and the typical direction of translocation for the molecules thought to be involved in CSF secretion are shown at the respective sites of expression.

Table 1 shows the Na^+^ transporting plasma membrane proteins reported to be expressed in the mammalian choroid plexus. Many of these proteins are Na^+^ extruders and are not considered further, while most others are carriers and channels not expected to take part in large-quantity Na^+^ uptake and not likely to be involved in transepithelial transport of a magnitude significant for CSF secretion. In the mouse, *slc4a10* encodes the Na^+^-HCO_3_^-^ import protein Ncbe (NBCn2 in humans), which is robustly expressed in the basolateral membrane of the CPECs along with the related Na^+^-HCO_3_^-^importer NBCn1 encoded by *slc4a7* (11, 12). Ncbe was suggested to mediate the DIDS (4,4′-Diisothiocyanatostilbene-2,2′-disulfonic acid) sensitive component and NBCn1 the DIDS-insensitive component of the Na^+^-dependent HCO_3_^-^ import in rat CP (12). Similar observations were obtained from mouse CPECs (13), however, the relative importance of the two transporters in Na^+^ transport and the relevance of other transporters remained elusive (Figure 1).

**Table 1.** Plasma membrane Na^+^ transporters of CPECs.

| Gene | Protein name | Protein abbreviation | Localization | Direction of Na<sup>+</sup> transport |
| --- | --- | --- | --- | --- |
| **Pumps** |  |  |  |  |
| *Atp1a1* | Sodium, potassium ATPase alpha 1 | Na/K-ATPse α1 | Luminal | Outward |
| *Atp1a2* | Sodium, potassium ATPase alpha 2 | Na/K-ATPse α2 | Luminal | Outward |
| *Atp1a4* | Sodium, potassium ATPase alpha 4 | Na/K-ATPse α4 | Luminal | Outward |
| *Atp1b1* | Sodium, potassium ATPase beta 1 | Na/K-ATPse β1 | Luminal | Outward |
| *Atp1b2* | Sodium, potassium ATPase beta 2 | Na/K-ATPse β2 | Luminal | Outward |
| *Atp1b3* | Sodium, potassium ATPase beta 3 | Na/K-ATPse β3 | Luminal | Outward |
| **Carriers** |  |  |  |  |
| *Slc4a5* | Sodium, bicarbonate cotransporter, electrogenic 2 | NBCe2 | Luminal | Outward |
| *Slc4a10* | Sodium-dependent chloride/bicarbonate exchanger | Ncbe | Basolateral | Inward |
| *Slc4a7* | Sodium, bicarbonate cotransporter, electroneutral 1 | NBCn1 | Basolateral | Inward |
| *Slc5a2* | Sodium, glucose transporter 2 | SGLT2 | Basolateral | Inward |
| *Slc5a3* | Sodium, myo-inositol transporter | Smit1 | Both | Inward |
| *Slc5a4* | Sodium, glucose transporter | SGLT3 | N/D | Inward |
| *Slc5a5* | Sodium, iodide transporter | NIS | N/D | Inward |
| *Slc5a6* | Sodium, biotin transporter | SMVT | N/D | Inward |
| *Slc6a8* | Sodium, creatin transporter | CRT | N/D | Inward |
| *Slc6a20a* | Sodium, proline transporter | Xt3s1 | N/D | Inward |
| *Slc12a2* | Sodium, potassium, 2 chloride cotransporter 1 | NKCC1 | Luminal | Both directions |
| *Slc13a5* | Sodium, citrate transporter | NaCT | N/D | Inward |
| *Slc13a4* | Sodium, sulfate transporter | NaS2 | N/D | Inward |
| *Slc17a2* | Sodium, urate transporter | NPT3 | N/D | Inward |
| *Slc20a2* | Sodium, phosphate transporter 2 | Pit-2 | Basolateral | Inward |
| *Slc38a1* | Sodium, amino acid transporter | SNAT1 | Basolateral | Inward |
| *Slc22a5* | Sodium, carnitine transporter | OCTN2 | N/D | Inward |
| *Slc22a17* | Sodium, organic cation transporter | BOCT | N/D | Inward |
| *Slc22a18* | Organic cation transporter protein 2 | ORCTL2 | N/D | Inward |
| *Slc23a2* | Sodium, ascorbic acid transporter | Svct2 | N/D | Inward |
| *Slc28a3* | Sodium, nucleoside transporter | Cnt3 | N/D | Inward |
| *Slc38a3* | Sodium, amino acid transporter | SNAT3 | Luminal | Inward |
| **Channels** |  |  |  |  |
| *Scn1a* | Voltage-gated sodium channel alpha 1 | Nav1.1 |  | Inward |
| *Scn8a* | Voltage-gated sodium channel | Nav1.6 |  | Inward |
| *Trpv4* | Transient receptor potential cation channel vallinoid 4 | TRPV4 | Luminal | Inward |

The current study was undertaken to determine the contribution of Ncbe to the Na^+^ uptake into murine CPECs. We assessed the Na^+^ import in isolated clusters of CPECs or intact tissue *ex vivo* with a Na^+^-sensitive reporter dye loaded into the cells. The experimental protocols included the removal and re-introduction of Na^+^ in salt solutions with and without the CO_2_/HCO_3_^-^ buffer system and analyzing the effects of known transport inhibitors as well as exploiting Ncbe-ko mice (14). Taken together, our results are consistent with Ncbe being the major basolateral Na^+^ importer of the CPECs in our experimental setup.

## Methods

### Animals

Male c57bl/6 mice (Taconic) of 8-26 weeks were used for the experiments. Where indicated, mice with and without *slc4a10* gene disruption (Ncbe-ko and Ncbe-wt, respectively) were obtained by heterozygous breeding and genotyping of the *slc4a10*-targeted knockout mouse model has been described previously (14). These mice were bred on c57bl/6 background, and both female and male mice aged 5–8 weeks were used for experiments. All procedures conformed to Danish animal welfare regulations. The authors are licensed to breed the mouse strain by The Animal Experiments Inspectorate, Ministry of Food, Agriculture, and Fisheries (j.n. 2012-15-2935-00004).

### Isolation of epithelial cells

Mice were euthanized under isoflurane anaesthesia, and CP tissues from all four ventricles of each animal were isolated in 4°C HEPES-buffered salt solution (HBS, pH 7.4, Table 2) under a stereomicroscope. Isolated CPs were digested into smaller cell clusters by 2 µg/ml dispase (Invitrogen) and 2 µg/ml collagenase B (Roche) in calcium-free HBS (Table 2) at 37°C for 30 min. Choroid plexus epithelial cells (CPECs) were washed and kept at 4°C until use within 5 hours between dissection and experiment.

**Table 2.**
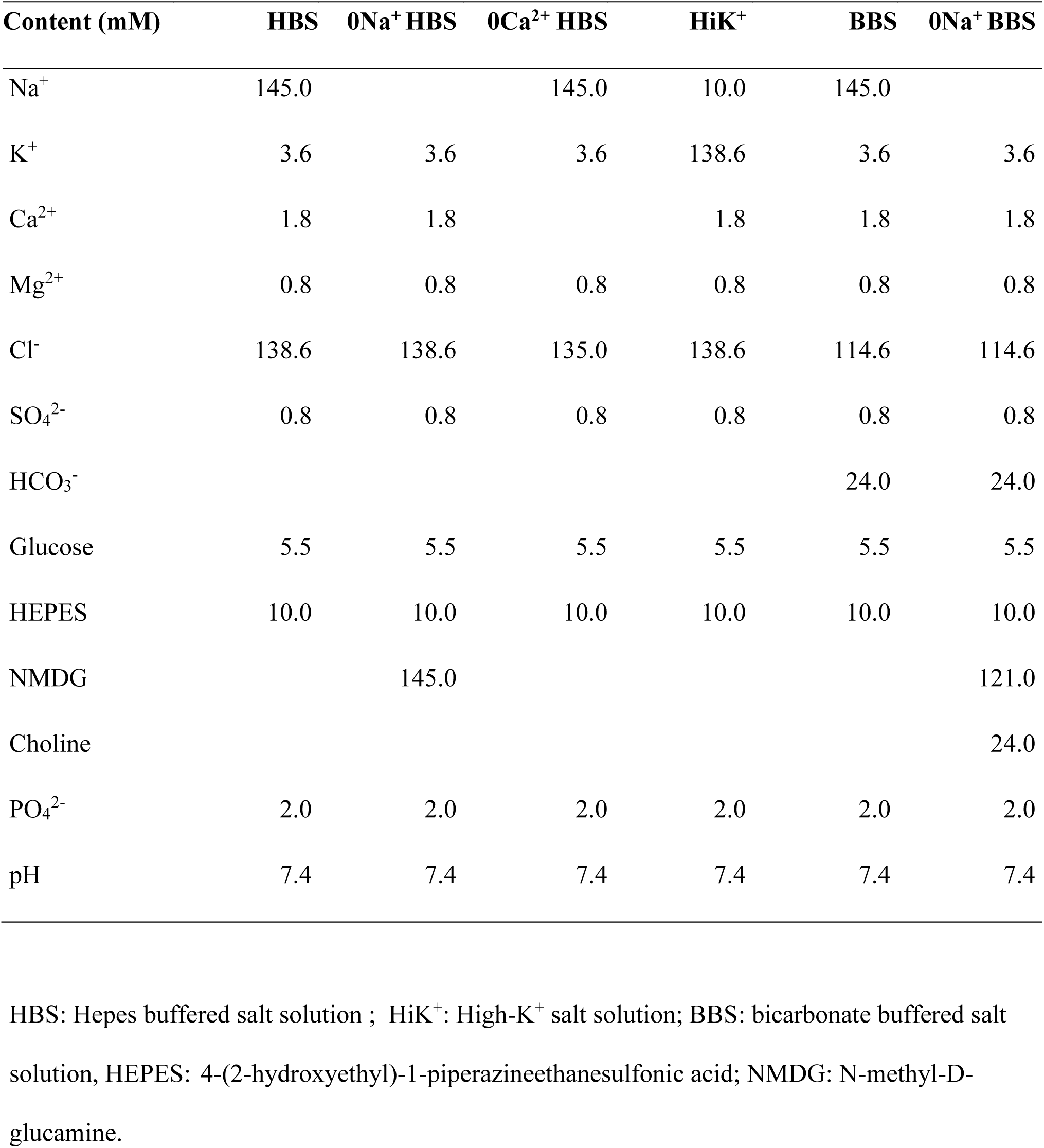
Experimental salt solutions.

| Content (mM) | HBS | 0Na<sup>+</sup> HBS | 0Ca<sup>2+</sup> HBS | HiK<sup>+</sup> | BBS | 0Na<sup>+</sup> BBS |
| --- | --- | --- | --- | --- | --- | --- |
| Na<sup>+</sup> | 145.0 |  | 145.0 | 10.0 | 145.0 |  |
| K<sup>+</sup> | 3.6 | 3.6 | 3.6 | 138.6 | 3.6 | 3.6 |
| Ca<sup>2+</sup> | 1.8 | 1.8 |  | 1.8 | 1.8 | 1.8 |
| Mg<sup>2+</sup> | 0.8 | 0.8 | 0.8 | 0.8 | 0.8 | 0.8 |
| Cl<sup>-</sup> | 138.6 | 138.6 | 135.0 | 138.6 | 114.6 | 114.6 |
| SO<sub>4</sub><sup>2-</sup> | 0.8 | 0.8 | 0.8 | 0.8 | 0.8 | 0.8 |
| HCO<sub>3</sub><sup>-</sup> |  |  |  |  | 24.0 | 24.0 |
| Glucose | 5.5 | 5.5 | 5.5 | 5.5 | 5.5 | 5.5 |
| HEPES | 10.0 | 10.0 | 10.0 | 10.0 | 10.0 | 10.0 |
| NMDG |  | 145.0 |  |  |  | 121.0 |
| Choline |  |  |  |  |  | 24.0 |
| PO<sub>4</sub><sup>2-</sup> | 2.0 | 2.0 | 2.0 | 2.0 | 2.0 | 2.0 |
| pH | 7.4 | 7.4 | 7.4 | 7.4 | 7.4 | 7.4 |

HBS: Hepes buffered salt solution ; HiK<sup>+</sup>: High-K<sup>+</sup> salt solution; BBS: bicarbonate buffered salt solution, HEPES: 4-(2-hydroxyethyl)-1-piperazineethanesulfonic acid; NMDG: N-methyl-D-glucamine.

### Intracellular Na^+^ and pH imaging

The cell clusters were mounted on Cell-Tak-coated coverslips (BD Biosciences) for 30 min at 37°C and loaded with the acetoxymethyl esters of either the intracellular Na^+^ sensitive probe Sodium Binding Fluorescent Indicator (SBFI-AM, 10 µM, 30 min., Invitrogen) or the intracellular pH (pHi)-sensitive probe 2’,7’-Bis-(2-Carboxyethyl)-5-(and-6)-Carboxyfluorescein (BCECF-AM, 2 µM, 5 min., Invitrogen). Coverslips were mounted in a closed perfusion chamber (RC-21BR; Harvard Apparatus) and superfused on the stage of an inverted iMic (Till Photonics) equipped with an Olympus UApo N340, 40x/1.35 NA oil immersion objective in a dark box constantly heated to 37°C. Till Vision software (Till Photonics) was used to control monochromator excitation with alternating wavelengths of 340 and 380 nm for SBFI, and 490 and 440nm for BCECF, exposure time (50 and 75 ms, respectively, for SBFI, 10 ms for both BCECF wavelengths), frequency (0.25 Hz), and binning (2×2 for SBFI). The light emission at 505–535 nm was recorded by a 14-bit cooled monochrome EMCCD camera (iXonEM+, Andor Technology) with 4.9× preamplifier gain (for SBFI) and 4×EM gain (both probes), and data were collected from user-defined ROIs of individual cells after background subtraction.

For SBFI, the single-cell fluorescence emission (I) ratio was recorded alternating at excitation 340 nm and 380 nm (I_340/380_). These I_340/380_ values were calibrated to [Na^+^]_i_ using Na^+^ ionophores in separate calibration experiments. SBFI calibration was performed in HBS at 0, 15, 50, and 145 mM Na^+^ with 10 µM of each of the ionophores gramicidin and monensin (Invitrogen). The empiric calibration curve was compared to a theoretical curve based on Ca^2+^ calibration by Grynkiewicz and coworkers (15), as SBFI is a derivative of 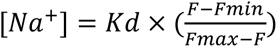, where Kd is the apparent dye Na^+^ dissociation constant, F is the variable observed fluorescence ratio, Fmin and Fmax are the fluorescence ratios at minimal (0 mM) and maximal (145 mM) [Na^+^]_o_, respectively. For most protocols, the [Na^+^]_i_ experiments were concluded by a 1- or 2-point calibration with 10 µM gramicidin to allow correction for day-to-day variation in the baseline I_340/380_. However, most attempts to obtain valid data from the end-point calibration failed because of massive cell swelling and blebbing upon gramicidin exposure. Instead, the I_340/380_ traces were normalized to the mean I_340/380_ values collected from all 45 individual experiments beginning with an HBS baseline recording. The rates of I_340/380_ recoveries, dI_340/380_/dt, were determined from the slopes of the traces as exemplified in the figures. The sequence of events was switched in protocols involving two pulses of Na^+^ removal and re-addition under different conditions (with/without CO_2_/HCO_3_^-^ or inhibitor exposure) to avoid unequal effects of exhaustion or photobleaching on the responses. The statistical analysis were performed with the normalized I_340/380_ values, and the apparent corresponding [Na^+^]_i_ was only computed from the mean I_340/380_ values and represents less accurate estimates.

For BCECF, the single-cell fluorescence emission was recorded at excitation 490 nm and 440 nm. The resulting I490/440 was calibrated to pH_i_ values using the ionophore nigericin (10 µM) in a high-K^+^ buffer at extracellular pH 6.2, 6.6, 7.0, 7.4, and 7.8. The intrinsic buffering capacity of mouse CPECs was determined previously and used to calculate the rates of net H^+^ recovery ([H^+^]i/dt) from the pH_i_ traces (16).

### Flame photometry

The choroid plexus was isolated from isoflurane-anaesthetized mice, weighed, and snap-frozen in liquid nitrogen for cell lysis, then 100 μL Milli-Q water was added, and finally boiled for 1 minute. The solubilized CPEC samples were diluted in 3 mM LiCl and processed for Na^+^ and K^+^ measurements by flame photometry (FLM3, Radiometer). Anaesthetized rats were placed on crushed ice, perfused for 5 min with 2-4°C isotonic PBS to cool down the animals and stop cellular transport processes, and then for 5 min with isotonic 2-4°C LiCl. The rat choroid plexus was isolated in isotonic LiCl at 4°C before removal of visible fluid and weighing of each choroid plexus tissue sample. Samples were handled as mouse samples from this point.

### Statistical analysis of functional data

All data are presented as scatter plots with mean values ± SD or simple mean bars. Two-tailed t-tests were generally used to compare changes between two groups in either paired or unpaired (Welch) analysis, as indicated (GraphPad Prism). Data from Ncbe-wt and -ko mouse CPECs were analyzed using a two-way ANOVA test comparing the mean values of three columns to a single reference column. A level of p<0.05 was considered adequate to indicate statistical significance. The exact p-values for mass spectrometry analysis are shown in each of the legends to Figures.

### Structure prediction

The sequence of the mouse NBCn2 protein (encoded by *Slc4a10*) was obtained from UniProt using accession Q5DTL9 (17). The full-length sequence contained 1,118 amino acid residues and was truncated at both the N- and C-termini. The truncation was based on a pairwise sequence alignment with the NDCBE protein (encoded by *SLC4A8*; PDB ID: 7RTM)(18) using the EMBOSS Needle tool on the EMBL website (19). The C-terminus was truncated by 66 amino acid residues, while the N-terminus was truncated by 481 amino acid residues. This resulted in a truncated sequence containing 571 amino acid residues per monomer. Because the predicted structure was a homodimer, the complete model contained 1,142 amino acid residues. The structure of the Ncbe protein was predicted using ColabFold (20, 21). The max-MSA depth was set to 512:1024, the num-recycle was set to 10, and the num-relax was set to 5 using Amber. The model type used was alphafold2_multimer_v3. The top-scoring (rank 1) structure was used as the starting structure. This protocol was adapted from that described by Johnsen *et al*. (22).

### System preparation

Two different protein dimer systems were prepared: Ncbe from the ColabFold prediction and NBCn1 (PDB ID: 9OVR (23). The two protein dimer systems were prepared using the Protein Preparation Workflow (24) in Schrödinger’s Maestro v2025.2. The protonation states of the amino acid residues were assigned using PROPKA at a pH of 7.4 (25, 26). Finally, the structures were minimized using the OPLS4 force field (27). This protocol was adapted from that described by Johnsen *et al*. (22).

### Molecular dynamics simulations

The Ncbe protein dimer was inserted into a membrane with a composition of 60% POPC and 40% cholesterol, and the box was solvated using OPC water. A salt concentration of 150 mM NaCl and 25 mM HCO₃⁻ was added to the solvent, and the system was neutralized using NaCl. System building was performed using CHARMM-GUI’s Membrane Builder (28, 29) by utilizing the CharmmGuiAuto.py script (30). The simulations were performed using the AMBER ff19SB force field (31) in GROMACS 2024.3 (32). The system was minimized using the steepest-descent algorithm for 10,000 steps. This was followed by four rounds of equilibration. First, an NVT simulation was performed with a time step of 1 fs for 0.25 ns. Second, an NPT simulation was performed with a time step of 1 fs for 0.25 ns. Third, an NPT simulation was performed with a time step of 2 fs for 0.5 ns. Fourth, an NPT simulation was performed with a time step of 2 fs for 10 ns. Restraints were applied to the backbone, side chains, dihedrals, and lipids during the equilibration steps, starting at 4,000, 2,000, 100, and 1,000 kJ/mol, respectively. The restraints were relaxed at each step, with only a 50 kJ/mol restraint on the backbone remaining during the final step.

The production runs were performed in the NPT ensemble at 310 K. Temperature was controlled using the V-rescale thermostat with a τt of 1.0 ps. Semi-isotropic pressure coupling was applied at 1 bar using the C-rescale barostat, with a τp of 5.0 ps and a compressibility of 4.5 × 10⁻⁵ bar⁻¹ (33, 34). The short-range interaction cutoff was set to 1.2 nm, and long-range electrostatic interactions were treated using the PME algorithm (35, 36). The van der Waals interaction cutoff was set to 1.2 nm, and the force-switch modifier was applied from 1.0 to 1.2 nm. The LINCS algorithm was applied to constrain bonds involving hydrogen atoms (37). The simulations were performed in triplicate, with each simulation lasting 500 ns and using a time step of 2 fs. This resulted in a total simulation time of 1.5 μs. The repeats were centered and fitted to the protein using the gmx trjconv tool (GROMACS manual). Clustering based on the root-mean-square deviation (RMSD) of the protein backbone was performed to obtain the medoid structure of the largest cluster. This structure was used for docking into Ncbe. Clustering was performed using the Python packages MDTraj, NumPy, and SciPy (38–40). This protocol was adapted from that described by Johnsen *et al*. (22).

### Ligand preparation

DIDS was built in Schrödinger’s Maestro v2025.2 and prepared using the LigPrep protocol. The protonation state was determined using Epik with a target pH of 7.4 ± 1.0 (41). DIDS was minimized using the OPLS4 force field (27).

### Induced Fit Docking

DIDS was docked using the extended-sampling protocol for Induced Fit Docking (IFD) in Schrödinger’s Maestro v2025.2 (42–44). DIDS was docked separately into both subunits of Ncbe and NBCn1. The box center was defined as the centroid of the following residues (Ncbe numbering): Thr569, Phe628, Ile632, Asp831, Val875, Ala876, Ala877, Thr878, and Lys1001. For Ncbe, this resulted in 57 DIDS poses in chain A and 39 DIDS poses in chain B. For NBCn1, this resulted in 33 DIDS poses in chain A and 49 DIDS poses in chain B.

### Covalent docking protocol

DIDS was covalently docked using the covalent docking protocol in Schrödinger’s Maestro v2025.2 (*45*). DIDS was docked separately into both subunits of Ncbe and NBCn1. Lys643 (Ncbe numbering) was used as the reaction site for cross-linking with DIDS.

### Molecular Mechanics - Generalized Born Surface Area

Prime Molecular Mechanics - Generalized Born Surface Area (MM-GBSA) rescoring of the DIDS poses was performed as a separate calculation for the IFD docking results and as part of the Covalent docking procedure in Schrödinger’s Maestro v2025.2. The protocol was performed using the OPLS4 force field (27) and the VSGB 2.1 implicit-solvent model (46). The default REAL-MIN protocol was used with receptor flexibility disabled. Missing side chains were not rebuilt, and energy decomposition was enabled. The membrane environment was not considered in the calculations.

### Analysis

The RMSD of the Ncbe backbone was calculated using gmx rms (GROMACS manual, (47)) (Supplemental Figure S7). The RMSF of Ncbe was calculated using gmx rmsf (GROMACS manual) (Supplemental Figure S8). The DIDS poses were clustered using RMSD based on the heavy atoms. This was performed using an approach similar to that used for the protein. Subsequently, all IFD poses were compared with the nearest CovDock pose using heavy-atom RMSD. A pose was classified as Covalent docking-like if the RMSD was at or below 3.5 Å. A pose was classified as productive if it satisfied the Covalent docking-like criterion and the distance between the Nζ atom of Lys643 and the nearest isothiocyanate carbon atom of DIDS was at or below 5.0 Å. This was done using MDTraj (38). Interactions between the proteins (Ncbe and NBCn1) and DIDS were calculated for all IFD poses using distance criteria. Interactions were identified using a custom geometry-based interaction-fingerprint script, with criteria adapted partly from the ProLIF defaults (48) and partly from predefined geometric heuristics.

### Summary statistics and visualizations

Plots were generated using Matplotlib, and structures were visualized using ChimeraX (49). Means and standard deviations were calculated using NumPy (39).

## Results

### CPEC SBFI signal and calibration to [Na^+^]_i_ values

It is well-known that obtaining useful SBFI signals from living cells can be challenging. Here we obtained suitable signal-to-noise ratio images by exploiting a highly sensitive camera and usual optimization of dye concentration and loading periods (Figure 2A). Although each calibration experiment yielded a clear dependence of the emitted SBFI fluorescence excitation ratio (I_340/380_) on [Na^+^] (Figure 2B), the inter-experimental variation in signal range was substantial, as illustrated by the calibration curve shown in Figure 2C. SBFI is modified from the state-of-the-art intracellular Ca^2+^ dye FURA2 and can also be calibrated according to the dye intensities at minimal and maximal ion concentrations. Thus, the observed calibration curve for SBFI closely resembles the curve calculated *ad modum* Grynkiewicz (15) in Figure 2C.

**Figure 2.**
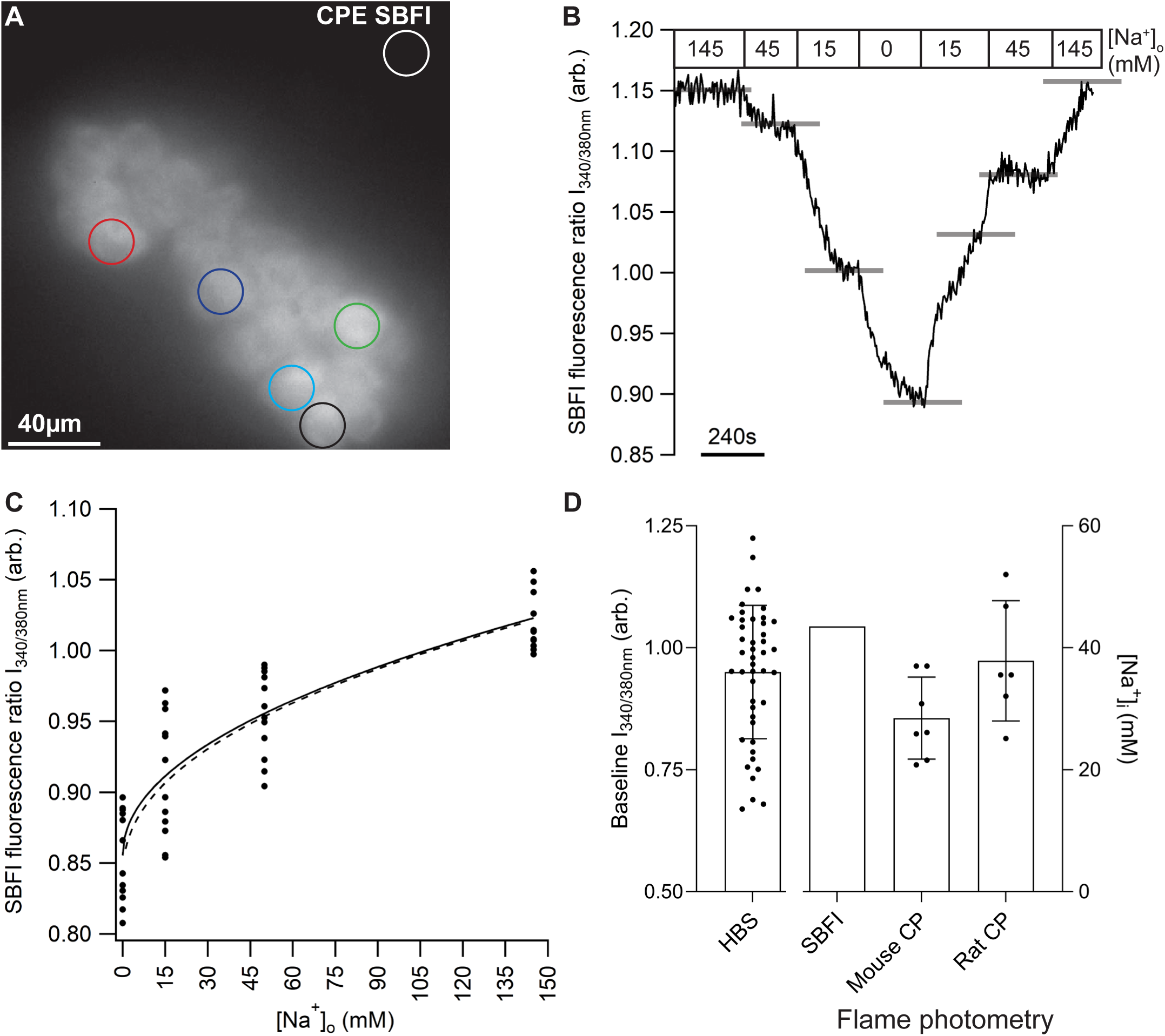
Calibration of the intracellular SBFI signal in isolated clusters of CPECs. A) A fluorescence micrograph of an SBFI-loaded cluster of CPECs from an Ncbe-wt mouse. The white circle indicates the background region; other colors exemplify cellular regions of interest. B) Representative trace of an SBFI calibration experiment. The fluorescence excitation ratio I_340/380_ is shown as a function of time. Boxes above the traces indicate the experimental conditions. The numbers 0-145 indicate the [Na^+^]_o_ during the experiment and gray lines mark the I_340/380_ level at the end of equilibration of [Na^+^]_i_ to [Na^+^]_o_ in gramicidin-permeabilized CPECs. C) The calibration curve of I_340/380_ as a function of [Na^+^]_o_ illustrated as a scatter plot (n=11), as well as the curve fit (power function, solid line) and the theoretical calibration curve (dotted line) calculated *ad modum* Grynkiewicz^5^: 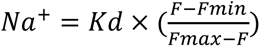 Bar graph comparing the baseline SBFI fluorescence ratio (scatter plot of all values obtained in HBS, bars indicate mean ± SD, n=45), the estimated [Na^+^]_i_ value corresponding to the mean baseline I_340/380_. The [Na^+^]_i_ value was obtained *ex vivo* by flame photometry of freshly isolated mouse choroid plexus or in rats after *in vivo* flushing out extracellular Na^+^ (scatter plot with bars indicating mean ± SD, n=7 for mice, n=6 for rats, respectively).

A variable baseline I_340/380_ was observed both between cells of a given experiment and between experiments (first bar, Figure 2D). We sought to correct for the large variation in baseline I_340/380_, by concluding each experiment with a 1- or 2-point calibration of [Na^+^] in the presence of gramicidin. This approach was abandoned, as most cells swelled and probably leaked dye during the procedure upon gramicidin exposure, rendering the signal-to-noise ratio worse than 2:1. Therefore, we have normalized each trace to the I_340/380_ at the end of the baseline period for the trace. We have only calibrated the observed mean I_340/380_ values to [Na^+^]_i_ levels (Figure 2D second bar). This estimated [Na^+^]_i_ > 40 mM from the 45 applicable experiments was higher than expected. We also determined [Na^+^]_i_ by flame photometry in mice and rats, where the extracellular Na^+^ was minimized by infusing the rats with ice-cold isotonic LiCl before isolating the choroid plexus for this analysis (left bars, Figure 2D). Taken together, we believe the method is technically sound for the analysis of Na^+^ transport in the choroid plexus - mainly as analyzed using the uncalibrated I_340/380_ and that the [Na^+^]_i_ was elevated compared to most transporting epithelia, at least in the *ex vivo* setting we applied.

### Approximately half the total Na^+^ uptake in CPECs depends on the presence of CO_2_/HCO_3_^-^

Figure 3A illustrates the import proteins believed to play major roles in Na^+^ transport in the CPE. The trace in Figure 3B shows the course of a typical recording of I_340/380_ when cells were first depleted of intracellular Na^+^ and then re-exposed to the ion, both in the presence or nominal absence of CO_2_/HCO_3_^-^. Recording of I_340/380_ was commenced after switching to Na^+^-free solution to reduce the total exposure time of the experiments and thereby the UV-induced photo-damage. The assessed minimal levels of I_340/380_ did not differ within the individual experiments (Figure 3C) and correspond to approximately 5 mM Na^+^. In general, the rates of I_340/380_-decreases during Na^+^ depletion seem slower than those of I_340/380_ increases upon re-addition of Na^+^, but this was outside the focus of the present study and, thus, not quantified. The systematically higher rate of I_340/380_ increase in the presence of CO_2_/HCO_3_^-^ compared to in their absence was statistically significant (Figure 3D). The CO_2_/HCO_3_^-^-dependent fraction amounted to ∼40% of the total I_340/380_ increase rate, corresponding to a ∼53% higher estimated mean [Na^+^]_i_ increase rate in the presence of CO_2_/HCO_3_^-^. The estimated [Na^+^]_i_ increase rates in the range 15-40 mM/min correspond well to the apparent time it took to return to baseline levels, as judged from the corresponding I_340/380_ traces.

**Figure 3.**
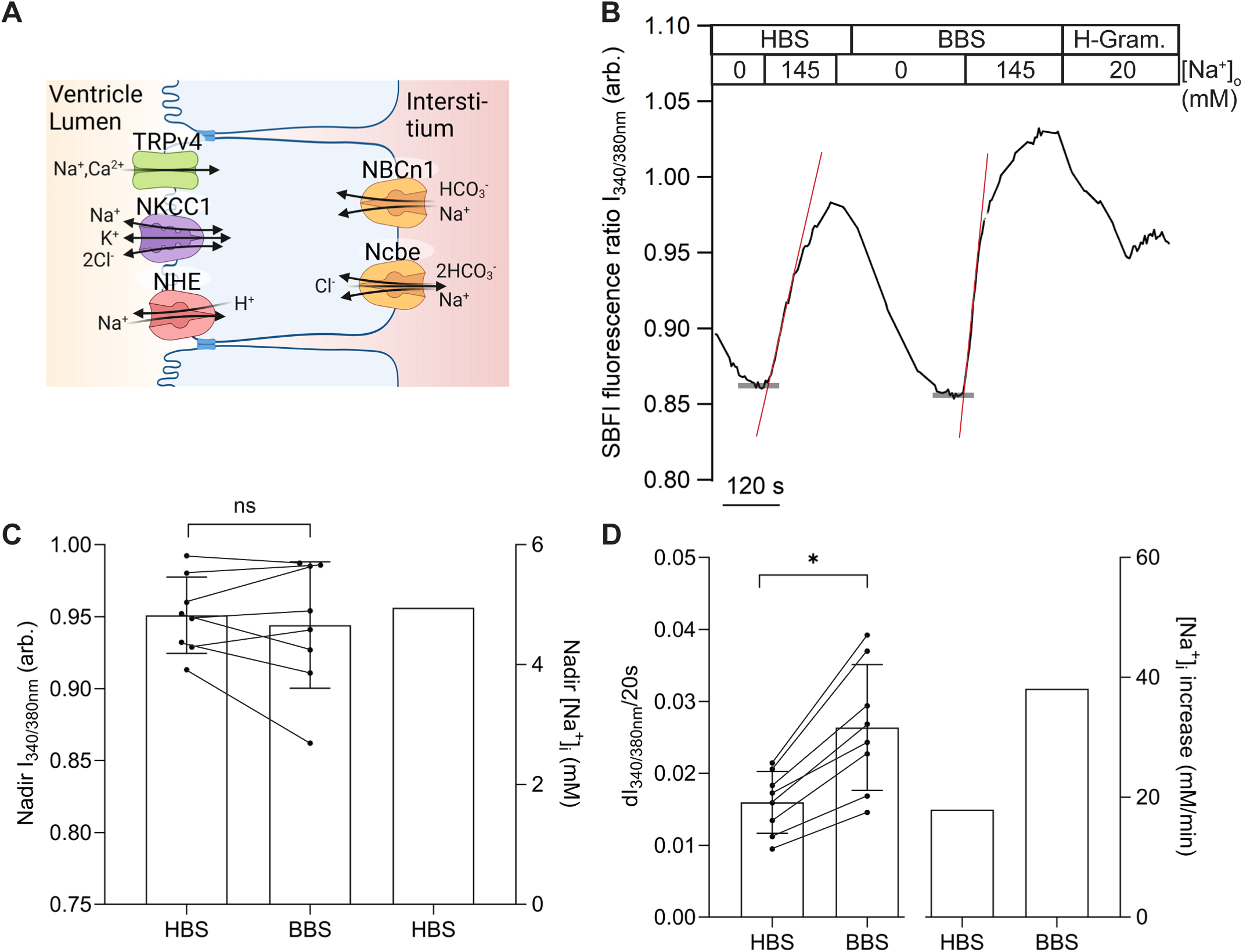
Comparison of the rates of SBFI fluorescence I_340/380_ increase in the presence and absence of CO_2_/HCO_3_^-^ in isolated clusters of CPECs. A) Schematic representation of sodium import pathways of a choroid plexus epithelial cell. B) Trace of SBFI fluorescence changes in isolated CPEC clusters in the course on an experiment. Boxes above the traces indicate the experimental conditions. Upper boxes indicate buffer type, where HBS is Hepes buffered Salt solution, and BBS is Bicarbonate buffered Salt solution. H-Gram indicates 10 µM gramicidin in HBS. The lower boxes indicate [Na^+^]_o_ in mM. Red lines indicate the slopes used to estimate the rate of change of I_340/380_, while thick gray lines mark the nadir I_340/380_. Grey sections of the trace indicate pauses. C) Scatter plot comparing the nadir SBFI fluorescence ratios obtained in the absence or presence of CO_2_/HCO_3_^-^ is shown on the left (n=8, p=0.4593, paired t-test. Bars indicate mean ± SD), while the bar on the right shows the corresponding [Na^+^]_i_ estimate in HBS. D) Scatter plot comparing the rates of SBFI fluorescence I_340/380_ increase in the presence and the nominal absence of CO_2_/HCO_3_^-^ is shown on the left (n=8), with bars indicating mean ± SD. The bars on the right show the corresponding estimates of [Na^+^]_i_ increase rates in HBS and BBS, respectively.

### The CO_2_/HCO_3_^-^-independent Na^+^ uptake takes place across the luminal membrane in CPECs

The results obtained by analyzing clusters of CPECs most likely reflect that both basolateral and luminal transporters contribute to the total Na^+^ depletion and recovery during our experiments. To assess the mechanisms of luminal Na^+^ recovery (Figure 4A), experiments were performed without enzymatic treatment of the choroid plexus (Figure 4B). Figure 4C illustrates the changes in I_340/380_ during experiments with Na^+^ depletion and re-administration in the isolated intact choroid plexus. Where indicated, 10 µM bumetanide was applied. The nadir I_340/380_ values obtained after Na^+^ removal are shown in Figure 4D and corresponded to an estimated [Na^+^]_i_ of 1.5 mM. Both the initial rate of I_340/380_ increase and the one just before/after bumetanide administration amounted to approximately 15 ms^-1^. The number corresponds well to the total rate of I_340/380_ increase observed in separated clusters of cells in the absence of CO_2_/HCO_3_^-^ in Figure 3D and the computed mean [Na^+^]_i_ increase rate of 17 mM/min under these conditions. Thus, the CO_2_/HCO_3_^-^-independent Na^+^ transport of the luminal membrane appears to equal that of the entire CPEC membrane.

**Figure 4.**
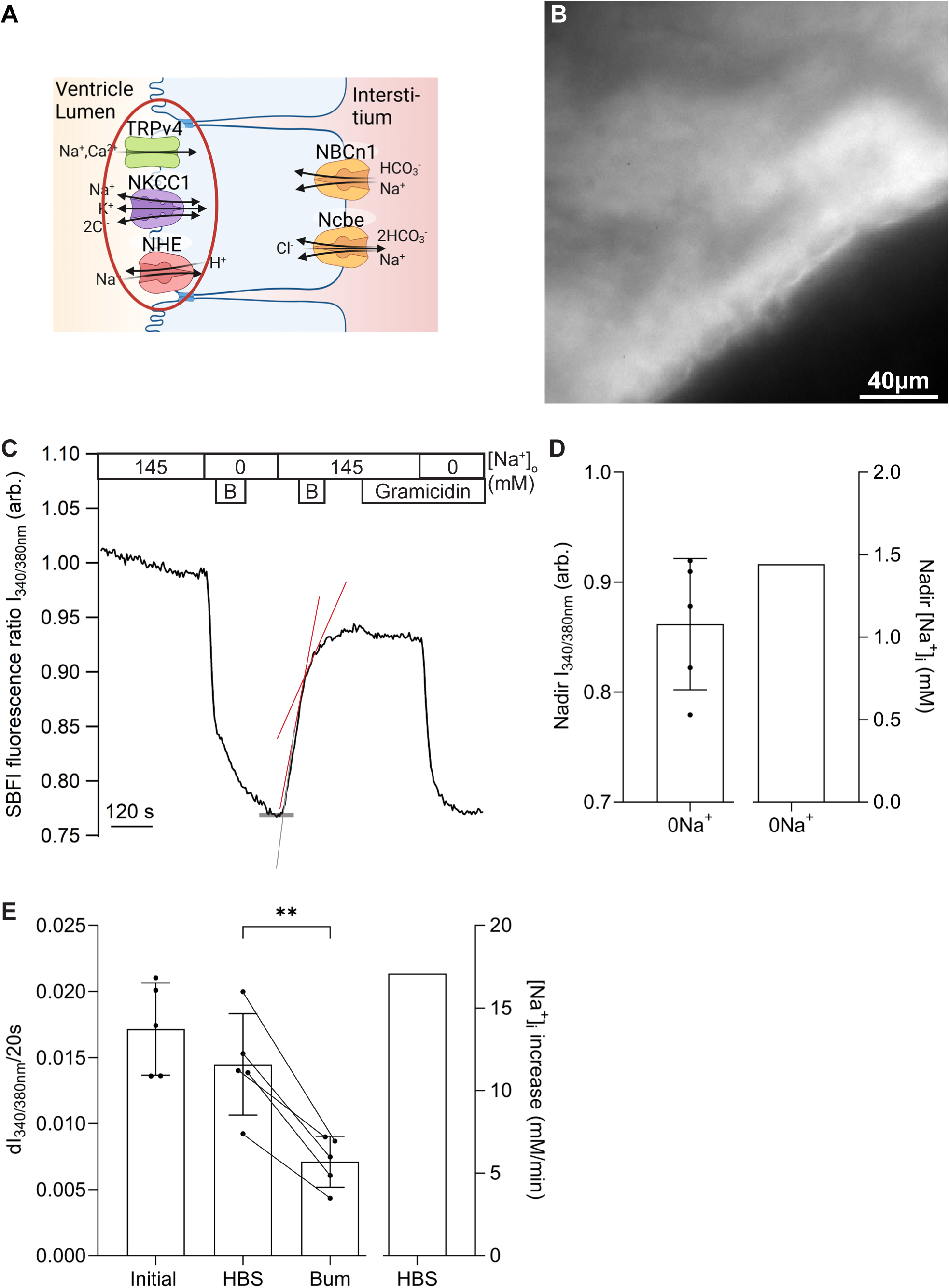
The rate of SBFI fluorescence I_340/380_ increase by Na^+^ import across the luminal membrane. A) Schematic representation of Na^+^ import pathways of a choroid plexus epithelial cell highlighting the luminal membrane importers. B) A fluorescence micrograph of an SBFI-loaded, undigested CP. C) Trace of SBFI fluorescence changes in undigested CPECs evoked by the experimental conditions. Upper boxes indicate [Na^+^]_o_ in mM, while lower boxes indicate periods with 10 µM bumetanide (B) or 10 µM gramicidin. Red lines indicate the slopes for measuring the rate of I_340/380_ change, while the initial slope of the I_340/380_ increase is indicated in grey. The thick gray line marks the nadir I_340/380_. D) Scatter plot of the nadir SBFI fluorescence ratios obtained in HBS is shown on the left (n=5, bars indicate mean ± SD), while the bar on the right shows the corresponding [Na^+^]_i_ estimate. E) Scatter plot showing the rate of SBFI fluorescence I_340/380_ increase at nadir I_340/380_ on the left and comparing the rate of SBFI fluorescence I_340/380_ increase with and without 10 µM bumetanide at the point of solution change (p=0.0059, paired t-test, n=5). Bars indicate mean ± SD. The bar on the right shows the corresponding estimate of the [Na^+^]_i_ increase rate in HBS.

### NKCC1 mediates ∼50% of the CO_2_/HCO_3_^-^-independent Na^+^ uptake in CPECs

In the undigested choroid plexus, administration of the NKCC1 inhibitor, Bumetanide, induced a statistically significant reduction of the I_340/380_ increase rate in paired analysis of the changes induced by addition or removal of the drug (Figure 4C and 4E). Figure 5A shows a cartoon of NKCC1 among the other Na^+^ import mechanisms of the CPECs.

**Figure 5.**
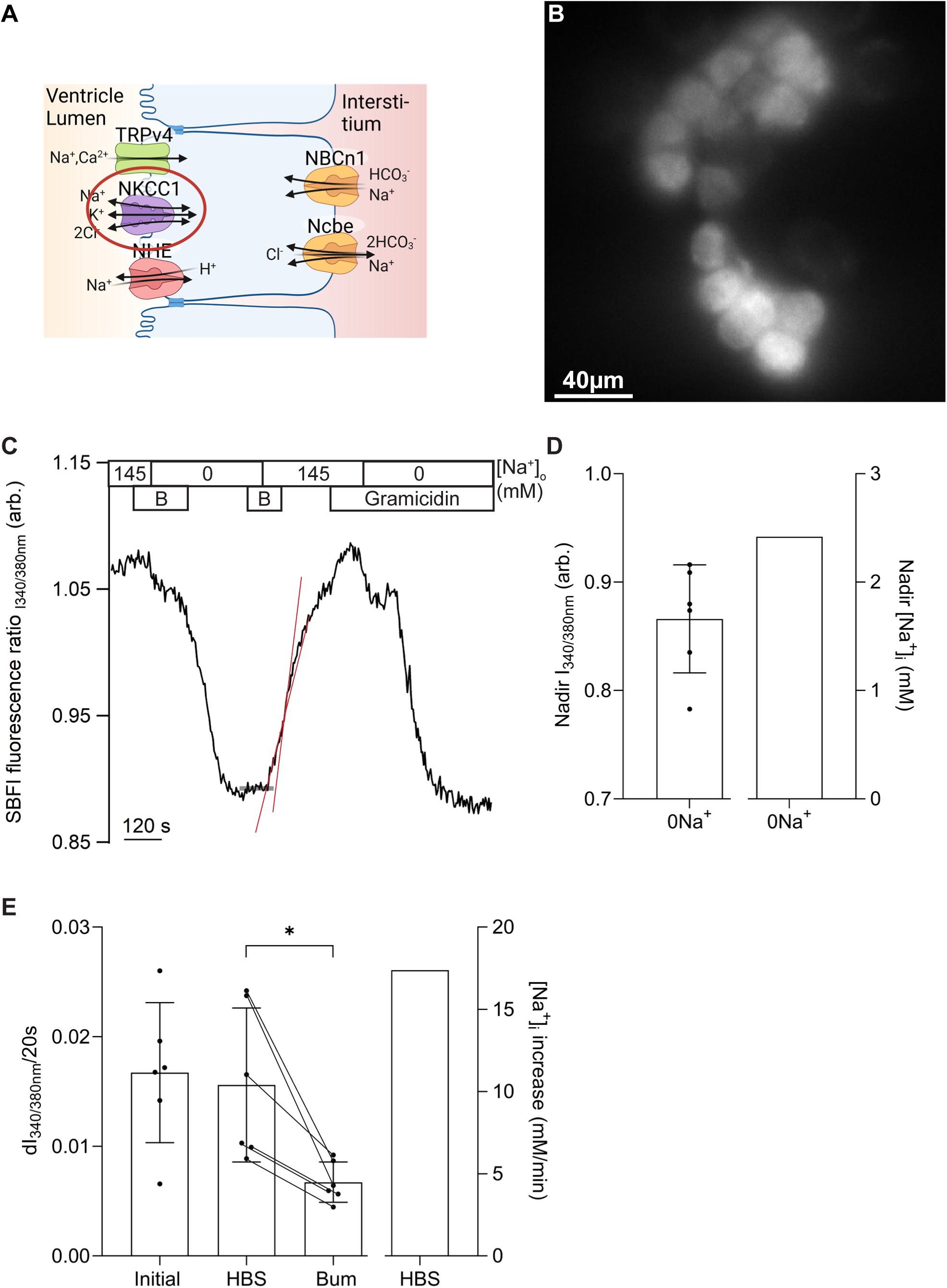
Effect of NKCC1 inhibitor bumetanide on the rate of SBFI fluorescence I_340/380_ increase in clusters of CPECs. A) Schematic representation of Na^+^ import pathways of a choroid plexus epithelial cell highlighting NKCC1 in the luminal membrane. B) A fluorescence micrograph of SBFI-loaded cluster of CPECs from a digested CP. C) Trace of SBFI fluorescence changes in the digested CPECs evoked by the experimental conditions. Upper boxes indicate [Na^+^]_o_ in mM, while lower boxes indicate periods with 10 µM bumetanide (B) or 10 µM gramicidin. Red lines indicate the slopes for measuring the rate of I_340/380_ change. The thick gray line marks the nadir I_340/380_. D) Scatter plot of the nadir SBFI fluorescence ratios obtained in HBS is shown on the left (n=6, bars indicate mean ± SD), while the bar on the right shows the corresponding [Na^+^]_i_ estimate. E) Scatter plot showing the rate of SBFI fluorescence I_340/380_ increase at nadir I_340/380_ on the left and comparing the rate of SBFI fluorescence I_340/380_ increase with and without 10 µM bumetanide at the point of solution change (p=0.0151, paired t-test; n=6). Bars indicate mean ± SD. The bar on the right shows the corresponding estimate of the [Na^+^]_i_ increase rate in HBS.

Isolated clusters of CPECs such as the one shown in Figure 5B were exposed to a protocol similar to the one applied to intact tissue. Figure 5C exemplifies the changes in I_340/380_ during experiments with Na^+^ depletion and re-administration in clusters of CPECs. Where indicated, 10 µM bumetanide was applied. The nadir I_340/380_ values obtained after Na^+^ removal are shown in Figure 5D and corresponded to an estimated [Na^+^]_i_ of 2.4 mM. Both the initial rate of I_340/380_ increase and the one just before/after bumetanide administration amounted to approximately 17 ms^-1^. Administration of Bumetanide to the CPEC clusters induced a statistically significant reduction of the rate of increase of I_340/380_ in paired analysis of the changes induced by addition or removal of the drug (Figure 5C and 5E).

### TRPv4 mediates 25-33% of the CO_2_/HCO_3_^-^-independent Na^+^ uptake in CPECs

Figure 6A shows a cartoon of TRPv4 among the other Na^+^ import mechanisms of the CPECs. In these experiments, the TRPv4 inhibitor RN1734 (30 µM) was applied 30 min before and throughout the experiment of Na^+^ depletion and re-administration (Figure 6B). Although there was a tendency towards higher values, RN1734 did not significantly alter the nadir I_340/380_, as illustrated in Figure 6C. The computed [Na^+^]_i_ corresponding to the mean nadir I_340/380_ was approximately 6 mM. As shown in Figure 6D, RN1734 significantly reduced the rate of I_340/380_ increase in paired analysis. The [Na^+^]_i_ recovery rate computed from the mean rate of I_340/380_ increases upon re-introduction of Na^+^ were approximately 22 mM/min (RN1734) and 29 mM/min (HBS). This amounts to approximately 33% of the control rate in CPECs judged by the I_340/380_ increase rates and 24% from the [Na^+^]_i_ recovery rate estimates (Figure 6D).

**Figure 6.**
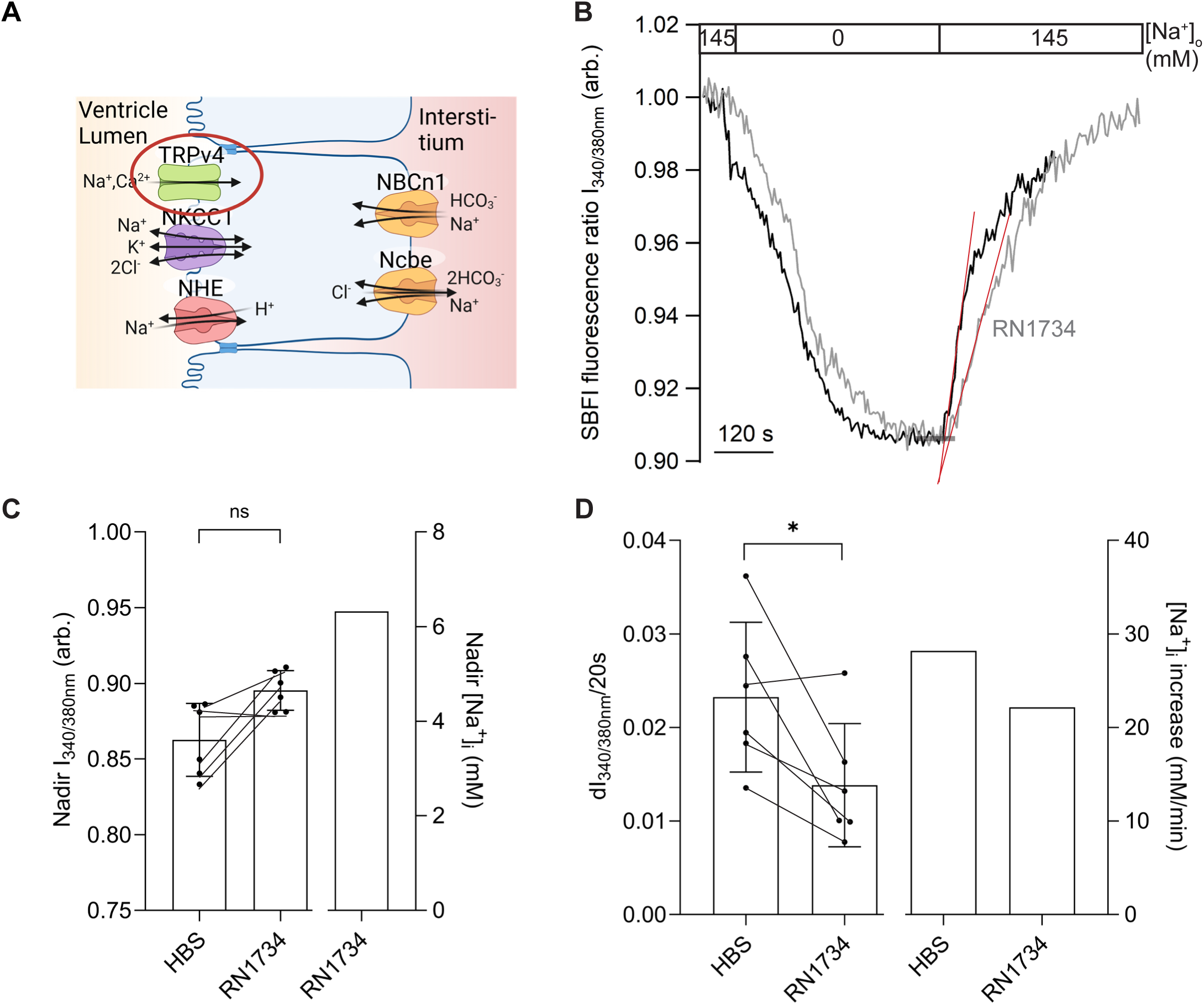
Effect of TRPv4 inhibitor RN1734 on the rate of SBFI fluorescence I_340/380_ increase in clusters of CPECs. A) Schematic representation of Na^+^ import pathways of a choroid plexus epithelial cell highlighting TRPv4 in the luminal membrane. B) Trace of SBFI fluorescence changes in the CPECs evoked by the experimental conditions. Boxes indicate [Na^+^]_o_ in mM. The black trace exemplifies the control experiments (HBS), while the grey trace represents a trace obtained in the presence of 30 µM RN1734 in HBS 30 min before the start and throughout the experiment. Red lines indicate the slopes for measuring the rate of I_340/380_ change. The thick gray line marks the nadir I_340/380_. C) Scatter plot shown on the left shows the nadir SBFI fluorescence ratios obtained in HBS and RN1734, respectively (p=0.1562, paired t-test; n=6). The bars indicate mean values ± SD, while the bar on the right shows the corresponding [Na^+^]_i_ estimate. D) Scatter plot on the left comparing the rates of SBFI fluorescence I_340/380_ increase at nadir I_340/380_ with and without 30 µM RN1734 (p=0.0352, paired t-test, n=6). Bars indicate mean values ± SD. The bars on the right show the corresponding estimates of the [Na^+^]_i_ increase rates in HBS and RN1734, respectively.

### NHEs mediate ∼1% of the CO_2_/HCO_3_^-^-independent Na^+^ uptake in CPECs

Figure 7A shows a cartoon of the Na^+^/H^+^ exchangers (NHE1 and NHE6) among the other Na^+^ import mechanisms of the CPECs. Amiloride, EIPA, benzamil and cariporide were first applied to assess the involvement of NHEs in the Na^+^ recovery of CPECs. Except for cariporide, all three inhibitors imposed massive, time-dependent buildup of cellular fluorescence in the presence of SBFI (not shown). Cariporide did not significantly affect baseline [Na^+^]_i_ or the [Na^+^]_i_ recovery rate (not shown), so the possible contribution of NHEs to CPEC Na^+^ transport was assessed by studying pH_i_ *in stead of* [Na^+^]_i_.

**Figure 7.**
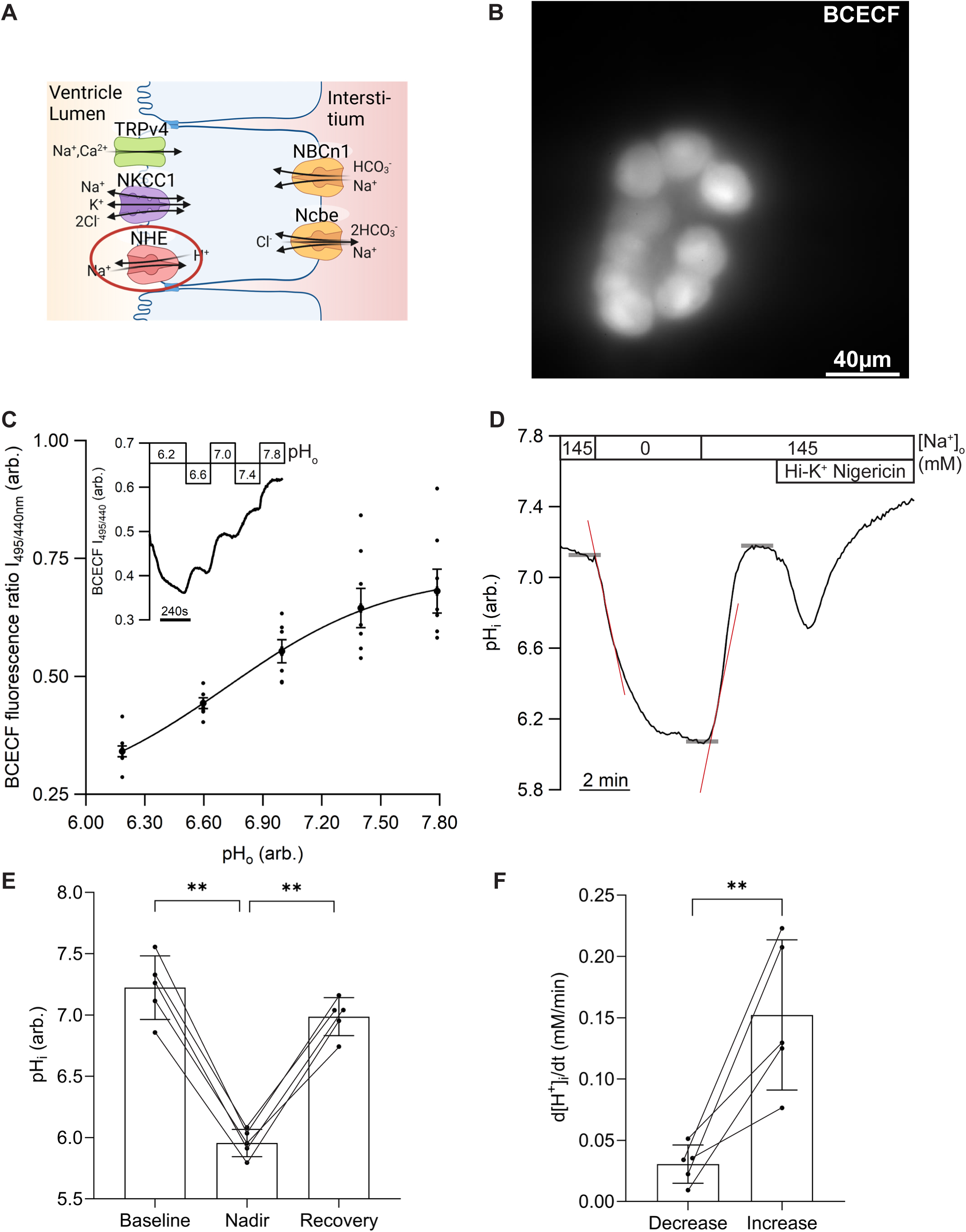
Changes in pH_i_ during [Na^+^]_i_ recovery of CPECs. A) Schematic representation of Na^+^ import pathways of a choroid plexus epithelial cell highlighting the NHE of the luminal membrane. Fluorescence micrograph of BCECF-loaded cluster of CPECs. C) The calibration curve of I_495/440_ as a function of pH_o_ illustrated as a scatter plot (n=7) with mean ± SD, as well as the polynomial curve fit (full line). The inset shows a representative trace of a BCECF calibration experiment. The fluorescence excitation ratio I_495/440_ is shown as a function of time. Numbers 6.2-7.8 indicate the pH_o_ during the experiment in permeabilized CPECs. D) Trace showing pH_i_ changes resulting from removal of and re-addition of Na^+^ to the perfusate and during 1-point calibration in nigericin-permeabilized cells (Hi-K/Nig pH 7.4). Upper boxes indicate [Na^+^]_o_ in mM, while the lower box indicates the calibration period (Hi-K Nigericin). Red lines indicate slopes at the points for measuring the rate of pH decrease and recovery upon removal or re-addition of Na^+^. The thick gray lines mark the baseline, the nadir, and the end of recovery pHi. E) Scatter plot of the individual values of initial, the nadir, and the end pHi, as indicated, during the experiment. Bars indicate mean values ± SD (Baseline to Nadir p=0.0004, nadir to Recovery p=0.0002, one-way ANOVA, n=5). F) Scatter plot illustrating the net H^+^-flux upon removal and re-addition of Na^+^ to the perfusate (p=0.0139, paired t-test; n=5). Bars indicate mean values ± SD.

Figure 7B shows a fluorescence micrograph of a BCECF-loaded CPEC cluster. This probe is known to penetrate into the nucleus from the cytosol, where the pH follows cytosolic pH. The calibration curve for converting the BCECF signal to pH_i_ values (Figure 7C) was based on experiments similar to the inset of the figure, where pH_i_ is clamped to pH_o_ of known values using nigericin as an ionophore. As expected, removal of extracellular Na^+^ was associated with a reduction of pHi, whereas Na^+^ re-introduction was associated with a fast recovery of pH_i_ (Figure 7D). The statistically significant changes in mean pH_i_ values before, during, and after the removal of Na^+^ from paired analysis are shown in Figure 7E. The mean [H^+^]i recovery rates during removal and re-introduction of extracellular Na^+^ were calculated from the rates of pH_i_ changes and the corresponding intrinsic buffering capacities at the given pH level and are summarized in Figure 7F. It is noted that the estimated [H^+^]i recovery rate is minimal compared to the corresponding [Na^+^]_i_ recovery rate: Given the 1Na^+^:1H^+^ stoichiometry of NHE1, the observed [H^+^]_i_ recovery rate of 0.15 mM/min amounts to less than 1% of the estimated [Na^+^]_i_ recovery rate. Thus, the NHEs seem to contribute insignificantly to Na^+^ transport in CPECs under the applied experimental conditions.

### Ncbe mediates most of the CO_2_/HCO_3_^-^-dependent and ∼ 50% of the total Na^+^ uptake in CPECs

Next we assessed the Na^+^ import of CPECs from Ncbe-knockout (-ko) mice and compared the results to those obtained with Ncbe-wildtype (-wt) littermate CPECs (shown in Figure 3) to consolidate the role of Ncbe as the main CO_2_/HCO_3_^-^-dependent Na^+^ uptake mechanism in these cells (Figure 8A). SBFI loading in Ncbe-ko CPECs was similar to Ncbe-wt CPECs with respect to signal intensity and the intracellular distribution that excludes the nucleus (Figure 8B). As in Figure 3B, the recording of I_340/380_ was started after switching to Na^+^-free solution to limit photo-damage (Figure 8C). The sequence of the elements of the protocol was also maintained in experiments of Ncbe-ko CPECs.

**Figure 8.**
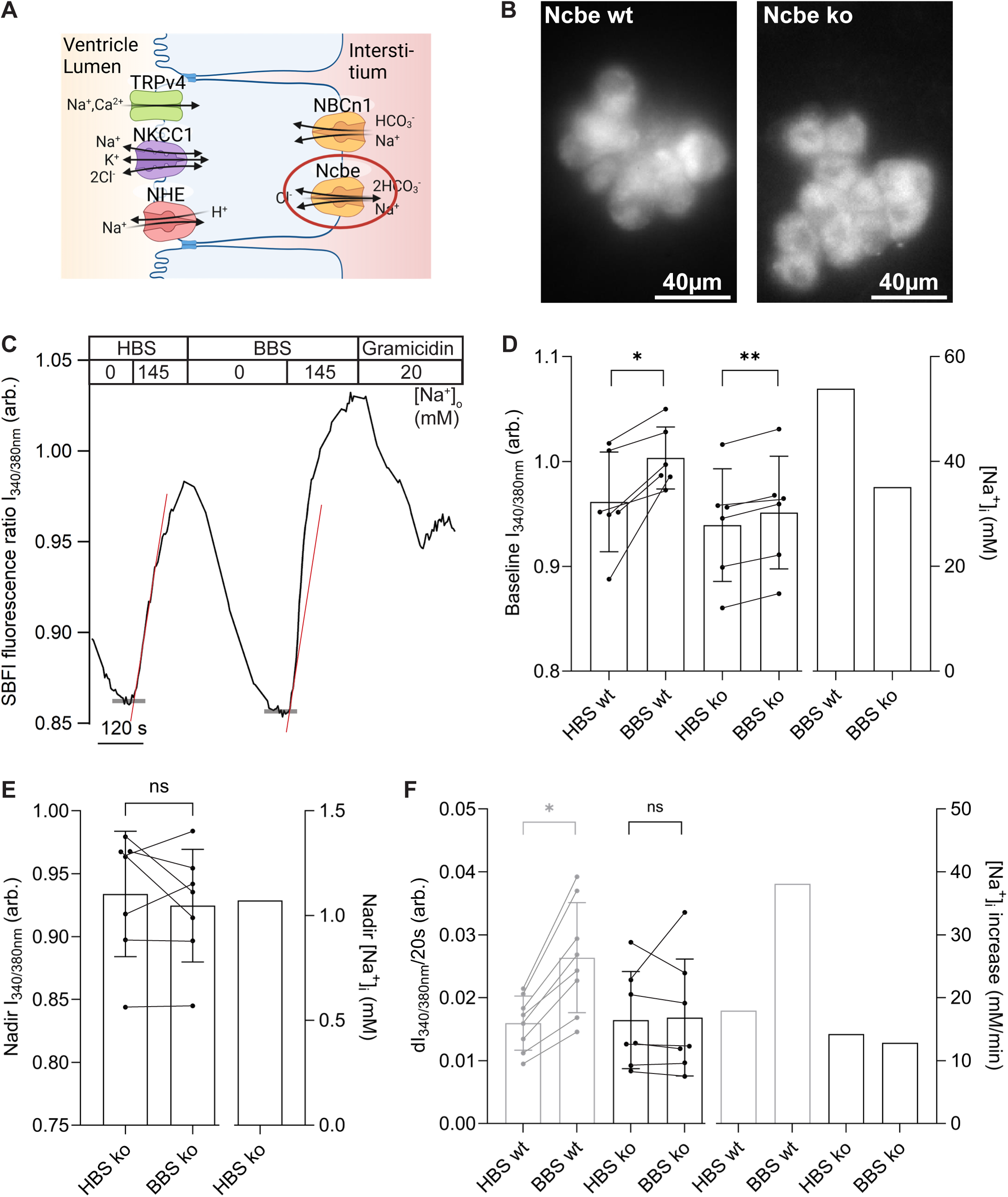
Comparison of the rates of SBFI fluorescence I_340/380_ increase in the presence and absence of CO_2_/HCO_3_^-^ in isolated clusters of CPECs from Ncbe-wt and -ko mice. A) Schematic representation of Na^+^ import pathways of a choroid plexus epithelial cell highlighting Ncbe in the basolateral membrane. B) Fluorescence micrographs comparing SBFI-loaded clusters of CPECs from Ncbe-wt and -ko mice. C) Trace of SBFI fluorescence changes in isolated CPEC clusters from an Ncbe-ko mouse. Boxes above the traces indicate the experimental conditions. Upper boxes indicate buffer type, where HBS is Hepes buffered Salt solution, and BBS is Bicarbonate buffered Salt solution. The lower boxes indicate [Na^+^]_o_ in mM. Red lines indicate the slopes used to estimate the rate of change of I_340/380_, while thick gray lines mark the nadir I_340/380_. D) Scatterplot comparing the baseline SBFI fluorescence ratios in HBS and BBS calculated for both Ncbe-wt mice and Ncbe-ko littermates. Bars indicate mean values ± SD. For Ncbe-wt mice, p=0.0143, while p=0.0001590 for Ncbe-ko mice (paired t-tests, n=6 for both genotypes). The bars on the right show the corresponding estimates of the [Na^+^]_i_ increase rates in BBS for Ncbe-wt and -ko mice, respectively. E) Scatter plot comparing the nadir SBFI fluorescence ratios obtained in the absence or presence of CO_2_/HCO_3_^-^ for Nbce-ko mice is shown on the left (n=7, p=0.4593, paired t-test, the bars indicate mean values ± SD), while the bar on the right shows the corresponding [Na^+^]_i_ estimate in HBS. F) Scatter plots comparing the rates of SBFI fluorescence I_340/380_ increase in the presence and the nominal absence of CO_2_/HCO_3_ ^-^ are shown on the left (n=7), with bars indicating mean values ± SD (n=7, p=0.8345, paired t-test). The bars on the right show the corresponding estimates of [Na^+^]_i_ increase rates in HBS and BBS, respectively. The Ncbe-wt data are from Figure 3 and indicated here in grey for comparison.

Baseline I_340/380_ was significantly higher in the presence of CO_2_/HCO_3_^-^ in both Ncbe-wt and - ko CPECs than without this buffer system in paired analysis (Figure 8D). There was a tendency toward a higher baseline I_340/380_ in Ncbe-wt than in Ncbe-ko cells (p= 0.0726 by unpaired t-test, as animals were not matched). The computed baseline [Na^+^] in the presence of CO_2_/HCO_3_^-^ was ∼54 mM for Ncbe-wt and *∼*35 mM for Ncbe-ko CPECs, respectively. Removal of extracellular Na^+^ yielded similar nadir I_340/380_ values in Ncbe-ko CPECs with and without CO_2_/HCO_3_^-^ (Figure 8E), with an estimated nadir [Na^+^]_i_ of ∼1 mM. As shown in Figure 8F, the mean I_340/380_ recovery rate was significantly higher in Ncbe-wt than in Ncbe-ko in the presence of CO_2_/HCO_3_^-^ buffer as was the I_340/380_ for both CPECs genotypes without the buffer system (p= 0.0273 by two-way ANOVA comparing Ncbe-wt BBS to the other columns). The computed [Na^+^]_i_ recovery rate in Ncbe-ko mice was in the range of 13-14 mM/min in both the presence and absence of CO_2_/HCO_3_^-^. Thus, the apparent Ncbe-dependent I_340/380_ recovery rate amounts to ∼ 36% of the total I_340/380_ recovery, corresponding to a Ncbe-dependence of 52% of the estimated total [Na^+^]_i_ recovery rate. Virtually all the CO_2_/HCO_3_^-^-dependent I_340/380_ recovery rate (94%) in Ncbe CPECs depends on Ncbe (Figure 8F).

### The HCO_3_^-^ transport inhibitor DIDS reduces the total Na^+^ uptake in CPECs by ∼50%

DIDS is a known inhibitor of most HCO_3_^-^ transporters and some Cl^-^ transporters and channels. In the context of CPEC Na^+^ transport, the drug is nevertheless useful, as it allows discrimination between the DIDS-sensitive Ncbe and the DIDS-insensitive NBCn1 (Figure 9A). The protocol of removing and re-introducing Na^+^ was adjusted compared to the previous experiments conducted in the presence of CO_2_/HCO_3_^-^ by the omission of this buffer system in the Na^+^-free buffer before the [Na^+^] recovery phase (Figure 9B). This was chosen to maximize the rate of Na^+^:HCO_3_^-^ transport during the recovery by securing an inward gradient for both transported species.

**Figure 9.**
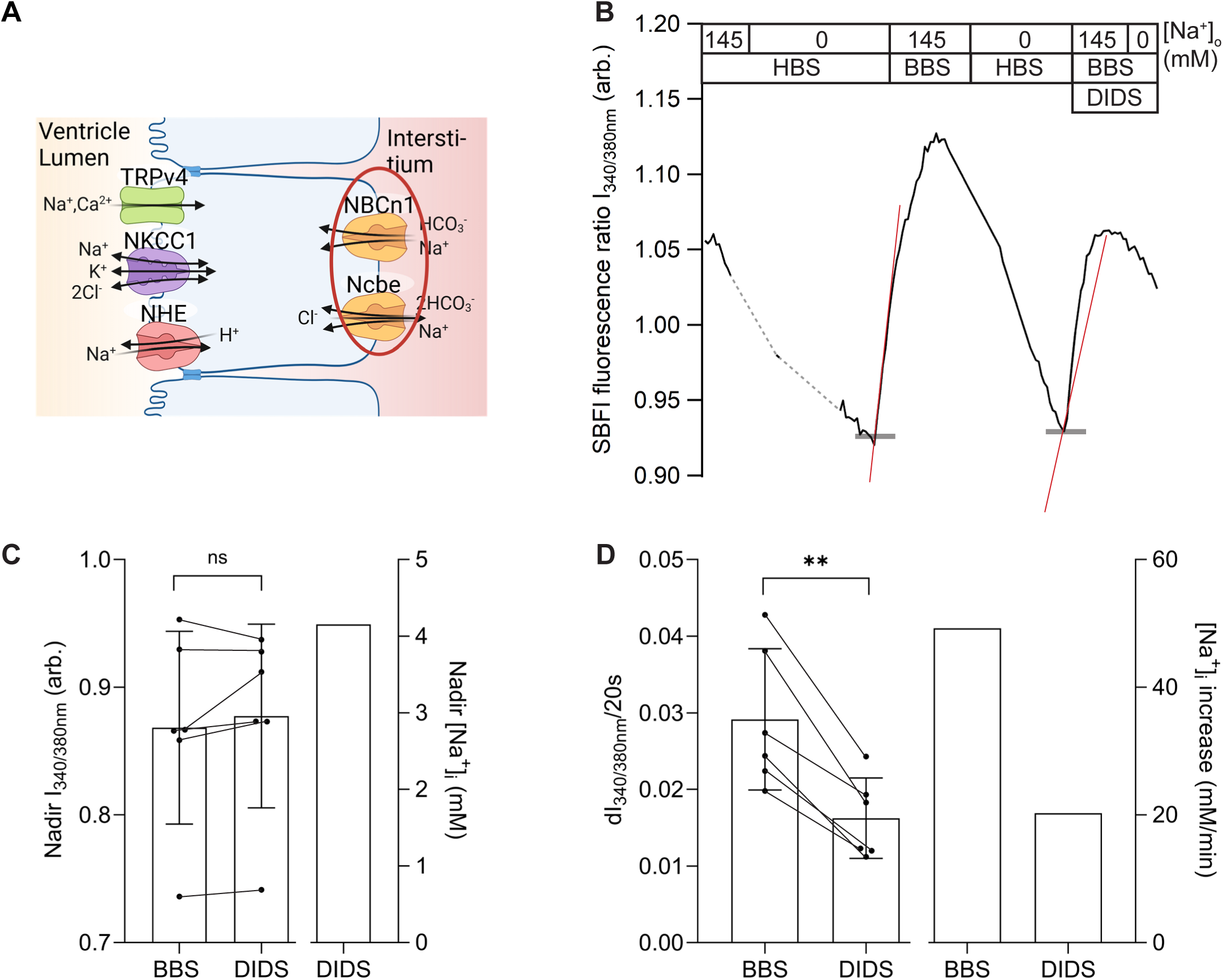
Effect of HCO_3_^-^ transport inhibitor DIDS on the rate of SBFI fluorescence I_340/380_ increase in clusters of CPECs. A) Schematic representation of Na^+^ import pathways of a choroid plexus epithelial cell highlighting the basolateral membrane transporters. B) Trace of SBFI fluorescence changes in the CPECs evoked by the experimental conditions. Upper boxes indicate [Na^+^]_o_ in mM. Middle boxes indicate buffer type, where HBS is Hepes buffered Salt solution, and BBS is Bicarbonate buffered Salt solution. The lower box marks the period of DIDS exposure (200 µM). Red lines indicate the slopes for measuring the rate of I_340/380_ change. The thick gray line marks the nadir I_340/380_. Grey sections of the trace indicate pauses. C) Scatter plot shown on the left shows the nadir SBFI fluorescence ratios obtained in HBS before exposure to BBS and BBS with DIDS, respectively (p=0.3239, paired t-test, n=6, bars indicate mean values ± SD), while the bar on the right shows the corresponding [Na^+^]_i_ estimate before DIDS exposure. D) Scatter plot comparing the rates of SBFI fluorescence I_340/380_ increase with and without 200 µM DIDS (p=0.0018, paired t-test, n=6). Bars indicate mean values ± SD. The bars on the right show the corresponding estimates of the [Na^+^]_i_ increase rates in BBS and BBS with DIDS, respectively.

As illustrated in Figure 9C, the large variation in nadir I_340/380_ appears to reveal higher day-to-day or inter-animal differences than the similar paired nadir values observed on a single day. Nevertheless, the mean nadir value yields a computed nadir [Na^+^]_i_ of approximately 4 mM. The rate of I_340/380_ increase after re-introducing Na^+^ was significantly inhibited by DIDS in paired analysis (Figure 9D). The I_340/380_ increase rate was reduced by 42%, which amounted to a 58% decrease in the corresponding estimated [Na^+^]_i_ recovery rate. Another set of experiments was conducted in the continued presence of CO_2_/HCO_3_^-^-throughout the Na^+^ challenges and yielded similar results. Here, the I_340/380_ increase rate was also reduced by 42% (n=6 controls, n=4 DIDS, p=0.0316 by unpaired t-test, Supplemental Figure S1). The corresponding estimated [Na^+^]_i_ recovery rates in these experiments were 46.2 and 21.1 mM/min for controls and DIDS-treated cells, respectively. Thus, the estimates of the DIDS-sensitive [Na^+^]_i_ recovery rate closely resemble the CO_2_/HCO_3_^-^-dependent [Na^+^]_i_ recovery rate of Figures 3-5, excluding a major role for NBCn1 in cellular Na^+^ uptake under the applied conditions.

### A putative structural background for differential action of DIDS on Ncbe and NBCn1

A computational chemistry approach was applied to assess whether DIDS was an appropriate drug to discriminate between Ncbe and NBCn1 transport. DIDS was docked into Ncbe and NBCn1 using Induced Fit Docking (IFD), which allowed local side-chain optimization, and analysis regarding putative covalent docking, in which DIDS was covalently linked to Lys643 of Ncbe (Figure 10A–C and Supplemental Figure S2). IFD generated 96 poses for Ncbe and 82 poses for NBCn1 across chains A and B. For Ncbe, the mean docking score was -7.1 kcal/mol, the mean IFD score was -32,299.2 kcal/mol, and the median MM-GBSA ΔG was -43.7 kcal/mol (Table S1). For NBCn1, the corresponding values were -7.4 kcal/mol, -40,936.0 kcal/mol, and -51.2 kcal/mol, respectively (Table S2). These results indicate that the docking procedure generated energetically favorable DIDS poses for both proteins. However, because the IFD score includes receptor-specific contributions, it was not used for direct comparison between NBCn1 and Ncbe. Covalent docking analysis generated two poses per chain for each protein. The mean affinity score and MM-GBSA ΔG were -3.8 and -20.5 kcal/mol, respectively, for Ncbe (Table S3) and -3.6 and -22.2 kcal/mol, respectively, for NBCn1 (Table S4).

**Figure 10.**
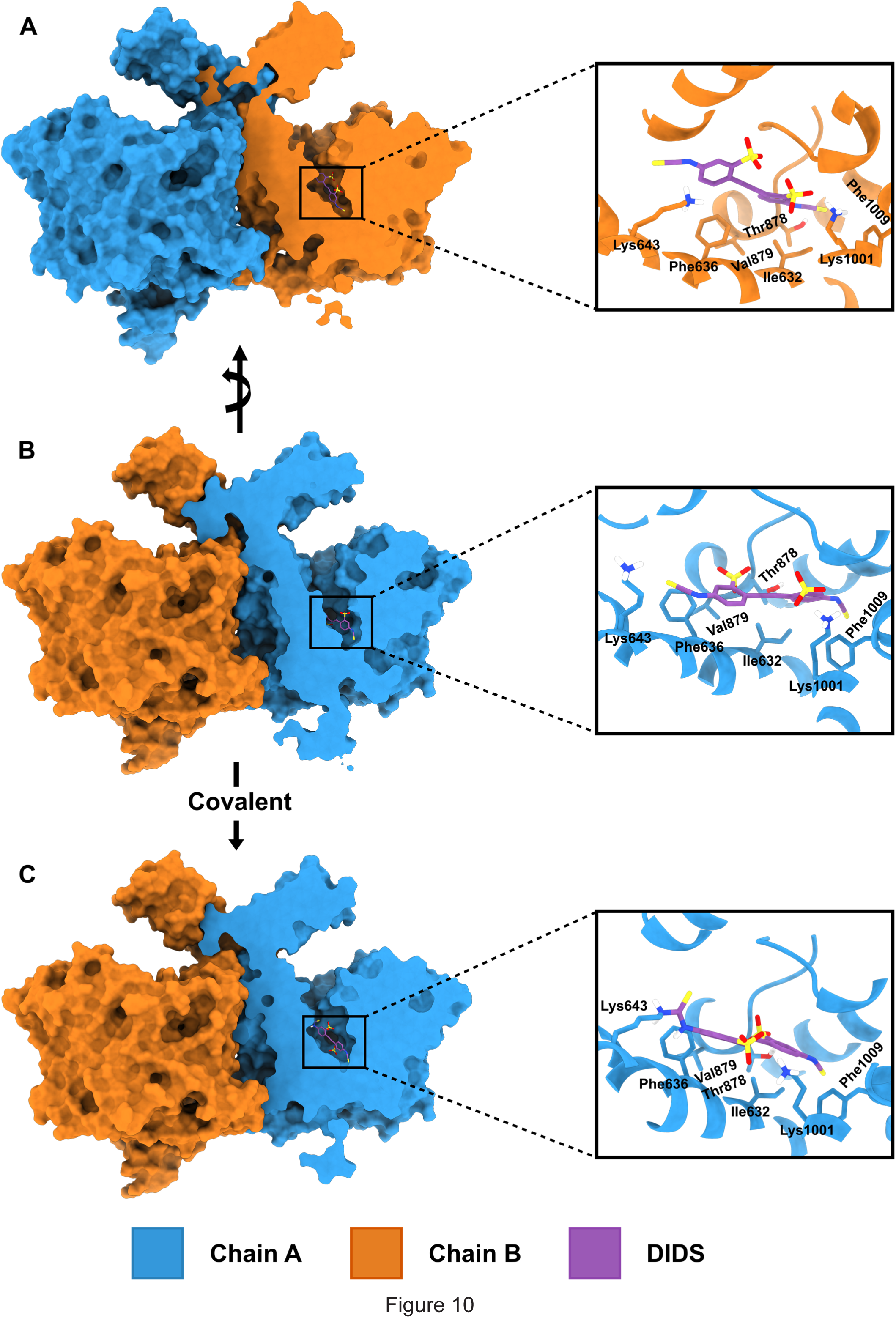
Docking poses for DIDS in the binding site of Ncbe. A) The sliced chain B subunit with DIDS docked using the IFD protocol. The zoom of the binding site shows the position and surrounding residues. B) Same as A), but for chain A subunit of Ncbe, where the structure has been rotated 180°. C) The sliced chain A subunit with DIDS docked using the Covalent docking protocol. The zoom of the binding site shows interacting residues and the covalently linked Lys643. Chain A is shown in blue, Chain B in orange, and DIDS in purple.

The IFD poses were divided into three groups: all IFD poses, Covalent docking-like poses with a heavy-atom RMSD of 3.5 Å or less from the nearest Covalent docking pose, and productive poses that additionally had a distance of 5.0 Å or less between the isothiocyanate carbon of DIDS and the Nζ atom of Lys643. For Ncbe, 46 of 96 poses (47.9%) were Covalent docking-like, and 25 poses (26.0%) were productive (Supplemental Figure S3 and S4). In comparison, 6 of 82 NBCn1 poses (7.3%) were Covalent docking-like, and 5 poses (6.1%) were productive (Supplemental Figure S3 and S4). In Ncbe, Ile632, Phe636, and Lys1001 had the highest interaction frequencies and remained prominent across the three pose groups (Supplemental Figure S5). The productive subset retained the Ile632/Phe636 hydrophobic region and the Lys1001 sulfonate anchor, while the participation of Lys643 and Thr878 increased. Additional contacts included Pro523, Ser568, Gly570, Ile635, Ala877, Val879, and Phe1009, although at lower frequencies. NBCn1 showed a broadly similar interaction pattern among all IFD poses (Supplemental Figure S6). However, Ile632, Phe636, and Lys1001 were not recovered in the smaller Covalent docking-like and productive subsets, which should be interpreted cautiously because these subsets contained only six and five poses, respectively. Thus, the comparative analysis of DIDS poses in Ncbe and NBCn1 models is consistent with – but less than a proof of - the drug having different effects on the two membrane proteins.

## Discussion

Na^+^ import across the basolateral membrane of CPECs is an absolute requirement for transcellular Na^+^ movement in CSF secretion. However, the immunolocalization and functional detection of luminal NHE1 activity in the mouse choroid plexus (13) rendered the molecular mechanisms by which Na^+^ is taken up by CPECs from the interstitial side elusive. In this study, we strived to identify the main Na^+^ import transport mechanisms of the mouse choroid plexus epithelium by assessing the Na^+^ changes induced by alternating buffer compositions regarding Na^+^ content, presence or absence of the CO_2_/HCO_3_^-^ buffering system, transport inhibitors, and by genetic knockout of a single candidate transporter.

Pioneering studies by Maren & Vogh and Johanson and coworkers exploited the radioisotope ^22^Na^+^ to directly assess Na^+^ concentrations, fluxes and their sensitivities towards various transporter and enzyme inhibitors (4–7, 50–52). Compared to most other transporting epithelia, the estimated [Na^+^]_i_ was relatively high, amounting to approximately 45 mM in rat CPECs *in vivo* by Johanson and coworkers (4, 5, 50). These estimates are the only reported *in vivo* data to date, to our knowledge, but correspond well to the [Na^+^]_i_ values obtained *in vitro* of 32 mM in rats (53), 31 mM in mice (54), and the values of the current study. A much lower estimate of 9 mM Na^+^ was recently reported by Gregoriades *et al.*, taking great care to keep the cells in cell medium at 37°C from isolation until experimentation (55). Both radiotracer and *in vitro* fluorometric approaches have inherent limitations, and the physiological [Na^+^] of CPECs therefore remains uncertain. SBFI provided internally consistent nadir values and rates of Na^+^ recovery in the present study, allowing us to assess the relative contributions of individual Na^+^ transport pathways with considerably higher temporal resolution than radiotracer approaches.

In the absence of a sided preparation of native mouse choroid plexus, dynamic [Na^+^]_i_ changes were determined in clusters of isolated CPECs. Consequently, we first assessed the contributions of known Na^+^ transporters irrespective of their subcellular localization. NKCC1, a highly expressed luminal transporter implicated in CSF secretion (51, 56–58) accounted for approximately 50% of the CO_2_/HCO_3_^-^-independent Na^+^ uptake under our experimental conditions. Although the physiological direction of NKCC1 transport remains debated (54, 55), is the transporter operates close to equilibrium (59) and may therefore mediate substantial Na^+^ entry when [Na^+^]_i_ is recovering far from steady state.

Unexpectedly, TRPv4 also contributed substantially to Na^+^ entry. TRPv4 is expressed in choroid plexus and conducts both Na^+^ and Ca^2+^ (60–62) but has not previously been considered a major Na^+^ import pathway (63, 64). Because temperature and extracellular osmolarity were kept constant, transient cell-volume changes during Na^+^ removal and re-addition may have promoted TRPV4 activation. By contrast, Na^+^/H^+^ exchange contributed only minimally to Na^+^ uptake, consistent with the relatively low abundance of NHE1 and NHE6 in choroid plexus (8, 10, 13).

Taken together, the results regarding the contributions of each of the transporters NKCC1, TRPv4 and the NHEs are important for interpretation of the observed total Na^+^ entry into CPRCs in the current study. The two significant contributors, NKCC1 and TRPv4, seem to account for approximately 75% of the CO_2_/HCO_3_^-^ independent Na^+^ transport (37% of the total Na^+^ import) in our experimental setup. However, they are all expressed exclusively in the CSF facing/luminal plasma membrane and are, thus, not relevant to the initial basic research question of what major pathways Na^+^ takes across the basolateral plasma membrane to sustain CSF secretion. The remaining approximately 25% of the CO_2_/HCO_3_^-^ independent Na^+^ transport must be ascribed to other Na^+^transporters, of which some are expressed in the basolateral membrane.

The remaining CO_2_/HCO_3_^-^-independent Na^+^ uptake may reflect activity of other basolateral Na^+^-dependent transporters, including phosphate, sulphate and glucose cotransporters, which were not examined individually in the present study. ENaC channel subunits have been identified in the choroid plexus by RT-PCR, immunoblotting, immunolabeling using both light and electron microscopy, and their regulation studied in disease models (65–68). The studies show expression mainly in the cytosol and plasma membrane expression with apparently higher luminal than basolateral membrane abundance. The ENaC-specific inhibitor, Benzamil, decreased [Na^+^]_i_ in the choroid plexus cells *in vitro* as a demonstration that the channel is a significant Na^+^ loader across the luminal membrane of the rat choroid plexus (67–69). However, we and others have repeatedly failed to demonstrate similar expressions of all ENaC subunits or benzamil sensitivity required for function in mouse CPECs (3, 10, 70). Given the discrepancy in findings between rats and mice and the unlikely predominantly cytosolic expression of ENaC even after experimental challenge to regulate abundance (53, 69), we tend to disregard ENaC as a major basolateral sodium entry pathway in the current study. The CO_2_/HCO_3_^-^-dependent contribution to the total Na^+^ re-entry to CPECs accounted for ∼50% of the total Na^+^ import with our experimental settings. Both electroneutral Na^+^:HCO_3_^-^ cotransporters NBCn1 and Ncbe are expressed in the basolateral membrane and could be implicated in the observed CO_2_/HCO_3_^-^-dependent fraction of the total Na^+^ re-entry as well as vectorial Na^+^ transport across the epithelium. Ncbe may be the first choice to consider further, as NBCn1 is occasionally expressed in the luminal membrane in some mouse strains and in parts of the human CP issue (13, 71). A significant contribution of the luminal membrane NBCe2 to the CO_2_/HCO_3_^-^-dependent Na^+^ uptake is unlikely, as it seems to only mediate outward transport in CPECs (72, 73). Indeed, the entire CO_2_/HCO_3_^-^-dependent fraction of the Na^+^ re-entry into CPECs was functionally absent in Ncbe-knockout mice, which strongly implicates Ncbe in the process. Importantly, the CPEC expression of the Na^+^ importers described above (CO_2_/HCO_3_^-^-dependent and CO_2_/HCO_3_^-^-independent) was unchanged by Ncbe knockout (74, 75) and therefore unlikely to be affected functionally by the genetic ablation. Direct evidence for the crucial involvement of Ncbe in CPEC Na^+^ transport was also articulated by the elevated baseline I348/380 in the presence of CO_2_/HCO_3_^-^. The observation indicates that Na^+^: HCO_3_^-^ import contributes to setting the steady-state [Na^+^]_i_ in isolated CPECs. This effect was numerically smaller in Ncbe-ko, suggesting that Ncbe is responsible for most of this phenomenon.

Bouzinova *et al.* demonstrated that the CO_2_/HCO_3_^-^ and Na^+^-dependent pH recovery from acidification in rat CPECs contained a DIDS-sensitive and a DIDS-insensitive component (12). The same phenomenon was later described in mouse CPECs (13). The findings were interpreted as evidence for activity of both Ncbe (DIDS sensitive) and NBCn1 (DIDS insensitive). Thus, while the pH recoveries suggest transport rates of similar magnitude for the two transporters, Na^+^:HCO_3_^-^cotransporters, the Na^+^ recovery in the present study was virtually all DIDS-sensitive using two different protocols of DIDS administration and was Ncbe-dependent. This apparent discrepancy between pH and Na^+^ fluorometry may be a product of the different experimental protocols applied. In pH recordings, we routinely remove Na^+^ and add CO_2_/HCO_3_^-^ simultaneously to obtain the lowest possible nadir pH and the maximal pH recovery rate. By contrast, the Na^+^ removal was conducted under CO_2_/HCO_3_^-^-free conditions and for longer periods to obtain lower nadir Na^+^, and therefore both pH_i_ and [Na^+^Ii would differ between the protocols. Another caveat to consider briefly is the fact that DIDS would also inhibit luminal NBCe2 and basolateral AE2. The latter does, however, not contribute to Na^+^ transport, and inhibition of outward transport by NBCe2 would tend to increase the apparent inward transport upon Na^+^ re-addition (73) and thereby tend to cause an underestimation of the contribution of Ncbe.

In interpreting DIDS inhibition of HCO_3_^-^ transporters, the research field relies on several point-mutational studies where specific amino acids were proven important for functional inhibition (76–78). The marked DIDS sensitivity of Ncbe compared with NBCn1 prompted us to examine whether structural differences could account for their differential inhibition. The conventional docking and MM-GBSA scores did not reveal an obvious difference that could explain the differential DIDS sensitivity of NBCn1 and Ncbe. Although some scores were slightly more favorable for NBCn1, the differences were small and did not consistently distinguish between the two proteins.

A clearer distinction emerged when the complete orientation of DIDS relative to the covalent docking products was considered. Covalent docking-like poses were 6.6-fold more frequent among the Ncbe docking poses than among the NBCn1 poses. When the additional Lys643 distance criterion was applied, productive poses were 4.3-fold more frequent for Ncbe. These fold differences describe the frequencies with which the docking workflow generated poses satisfying the geometric criteria and should not be interpreted as equilibrium probabilities. The results therefore suggest that both NBCn1 and Ncbe can accommodate generic DIDS binding, but that Ncbe more readily adopts orientations compatible with the proposed covalent reaction. In particular, Ncbe could simultaneously maintain interactions with the Ile632/Phe636 hydrophobic region and the Lys1001 sulfonate anchor while positioning the reactive group of DIDS near Lys643. The experimentally observed difference in DIDS inhibition may therefore reflect the probability of forming a productive pre-reaction complex rather than the stability of a generic bound state.

The ion binding-site sequence is highly conserved between NBCn1 and Ncbe and does not identify a single residue that can readily explain the difference in DIDS sensitivity. Instead, the distinction may arise from subtle differences in binding-site geometry, side-chain organization, pocket dimensions, or conformational dynamics. The docking results showed substantial variation between the two chains of Ncbe, indicating that productive docking is highly conformation-dependent and may be influenced by the preceding molecular dynamics simulations. Furthermore, Ncbe was represented by a medoid structure obtained from molecular dynamics, whereas NBCn1 was prepared directly from an experimental PDB structure. Differences between the proteins may therefore partly reflect their starting conformations and preparation procedures.

The interaction fingerprints were calculated from static poses using geometric criteria and did not establish the persistence of individual contacts. The MM-GBSA calculations also omitted the membrane environment, which may influence the energetics of ligand binding in these membrane proteins. Finally, the proposed covalent modification of Ncbe at Lys643 is based on DIDS-bound structures of AE1, AE2, and AE3 (79–81) but has not, to our knowledge, been demonstrated experimentally for Ncbe. The present results should therefore be considered a mechanistic hypothesis that requires validation through molecular dynamics simulations and experimental studies.

Several lines of evidence were presented implicating Ncbe in a major fraction of the Na^+^ uptake in mouse CPECs in this work. The high transport rate compared to other Na^+^ uptake pathways, together with its obligate expression in the basolateral membrane, makes Ncbe an attractive candidate to mediate Na^+^ entry from the interstitial side of the CPECs to sustain CSF secretion. One should acknowledge that the estimated Na^+^ transport rates with the current protocol *in vitro* may not precisely reflect their contribution to CSF secretion at steady state *in vivo*. However, the significance of Ncbe for setting the baseline [Na^+^]_i_ indicates a Ncbe-mediated transport activity close to *in vitro* steady state. A major role for Ncbe in basolateral Na^+^ import or CSF secretion by CPECs is also consistent with most of the early studies showing HCO_3_^-^-dependence, DIDS sensitivity, and even acetazolamide sensitivity (4–7, 50–52). The latter would reflect a case where membrane-bound carbonic anhydrases are promoting CO_3_^2-^ as a substrate for transport through Ncbe instead of HCO_3_^-^. Based on our present findings, we suggest Ncbe to be the most attractive future target to obtain pharmaceutical CSF inhibition. Also, in contrast to Ncbe expressed elsewhere in the CNS, Ncbe of the CPECs has the advantage of being directly exposed to the blood through fenestrated capillaries. We find that the suggested Ncbe-selective inhibition by DIDS over NBCn1 offers hope for designing such future therapeutics. Although DIDS itself is not a useful drug, it seems that the subtle differences in ion-binding pockets

In conclusion, the luminal Na^+^ uptake amounts to ∼ 50% of the total Na^+^ uptake of isolated choroid plexus epithelial cells. NKCC1 contributes ∼50% and TRPv4 contributes 25% of the luminal Na^+^ uptake upon re-addition of extracellular [Na^+^]. Na^+^/H^+^ exchange activity apparently contributes less than 1% of the Na^+^ uptake in isolated choroid plexus cells by our protocol. DIDS inhibited ∼ 50% of the total Na^+^ uptake, indicating that Ncbe contributes more than NBCn1 to CPEC Na^+^ import.

The docking analysis presented in this study indicated that DIDS could bind within the ion-binding sites of both NBCn1 and Ncbe. However, DIDS sampled a greater proportion of poses resembling the covalently linked pose in Ncbe than in NBCn1. The overall binding-site residues are highly conserved between the two proteins and do not immediately indicate why Ncbe is inhibited by DIDS while NBCn1 is less sensitive to DIDS. Instead, the variation between Ncbe and NBCn1 may arise from differences in the overall binding-site geometry, side-chain organization, and conformational dynamics. However, docking and MM-GBSA calculations alone cannot establish differential inhibition or covalent modification, and these findings should therefore be considered hypothesis-generating. Ncbe is the dominant basolateral membrane Na^+^ uptake mechanism of the choroid plexus epithelial cells and seems to contribute to setting baseline [Na^+^]_i_ *ex vivo*. Thus, Ncbe is a strong candidate as the rate-limiting step in CSF secretion.

## Acknowledgements

Nicoline Beiskjær, Mads Vaarby Sørensen, Peter Aakær Nielsen, and Kien Strand Olesen are thanked for expert technical assistance.

## Supplemental Material

**Supplemental Table S1.** Ncbe – DIDS IFD results. The docking score, IFD score, and dG value for all clusters in both chains A and B for Ncbe.

**Supplemental Table S2.** NBCn1 – DIDS IFD results. The docking score, IFD score, and dG value for all clusters in both chain A and B for NBCn1.

**Supplemental Table S3.** Ncbe – DIDS CovDock results. The CovDock scores and dG values for both poses in each chain for Ncbe.

**Supplemental Table S4.** NBCn1 – DIDS CovDock results. The CovDock scores and dG values for both poses in each chain for NBCn1.

**Supplemental Figure S1.** Effect of the HCO_3_^-^ transport inhibitor DIDS on the rate of SBFI fluorescence I_340/380_-increase in clusters of CPECs. Compared to Figure 9, the protocol was executed in the continued presence of CO_2_/HCO_3_^-^, and only one Na^+^-free challenge was induced per experiment. The scatter plot comparing the rates of SBFI fluorescence I_340/380_ increase with and without 200 µM DIDS (p=0.0316, unpaired t-test, n=6 for controls, n=4 for DIDS-treated cells, respectively). Bars indicate mean values ± SD. The bars on the right show the corresponding estimates of the [Na^+^]_i_ increase rates in BBS and BBS with DIDS, respectively.

**Supplemental Figure S2.** Docking poses for DIDS in the binding site of NBCn1. A) The sliced chain B subunit with DIDS docked using the IFD protocol. The zoom of the binding site shows the position and surrounding residues. B) Same as A), but for chain A subunit of NBCn1, where the structure has been rotated 180°. C) The sliced chain A subunit with DIDS docked using the Covalent docking protocol. The zoom of the binding site shows interacting residues and the covalently linked Lys643. Chain A is shown in blue, Chain B in orange, and DIDS in purple.

**Supplemental Figure S3.** RMSD of IFD poses to nearest CovDock pose. The RMSD was calculated for IFD poses in each chain for both Ncbe and NBCn1. The cutoff for being CovDock-like was set to an RMSD value below 3.5 Å (dashed line). Chain A is shown in blue and chain B in orange.

**Supplemental Figure S4.** The population of CovDock-like and productive poses. The percentages of IFD poses being CovDock-like (solid bars) and productive poses (shaded bars) are shown individually for each chain for both Ncbe and NBCn1. Productive poses are the subset of the CovDock-like poses that also are with the Nζ atom of Lys643 and the nearest isothiocyanate carbon atom of DIDS was at or below 5.0 Å. Chain A is shown in blue and chain B in orange.

**Supplemental Figure S5.** Interaction frequency between Ncbe and DIDS. Here the subsets of IFD poses are shown and the percentage of the population interacting with the specific residue. All IFD poses are shown in yellow, CovDock-like poses in green, and productive poses in purple.

**Supplemental Figure S6.** Interaction frequency between NBCn1 and DIDS. Here the subsets of IFD poses are shown and the percentage of the population interacting with the specific residue. All IFD poses are shown in yellow, CovDock-like poses in green, and productive poses in purple.

**Supplemental Figure S7.** Ncbe protein RMSD. The RMSD of the protein backbone during the simulations for each chain and all three repeats. Chain A is shown in blue and chain B in orange.

**Supplemental Figure S8.** Ncbe protein RMSF. The time averaged RMSF of the protein residues during the simulations for each chain and all three repeats. Chain A is shown in blue and chain B in orange. The transmembrane helices of Ncbe are shown as grey bars.

