## Supplementary material for "Ncbe is the main basolateral Na^+^ loading mechanism of the choroid plexus epithelium"

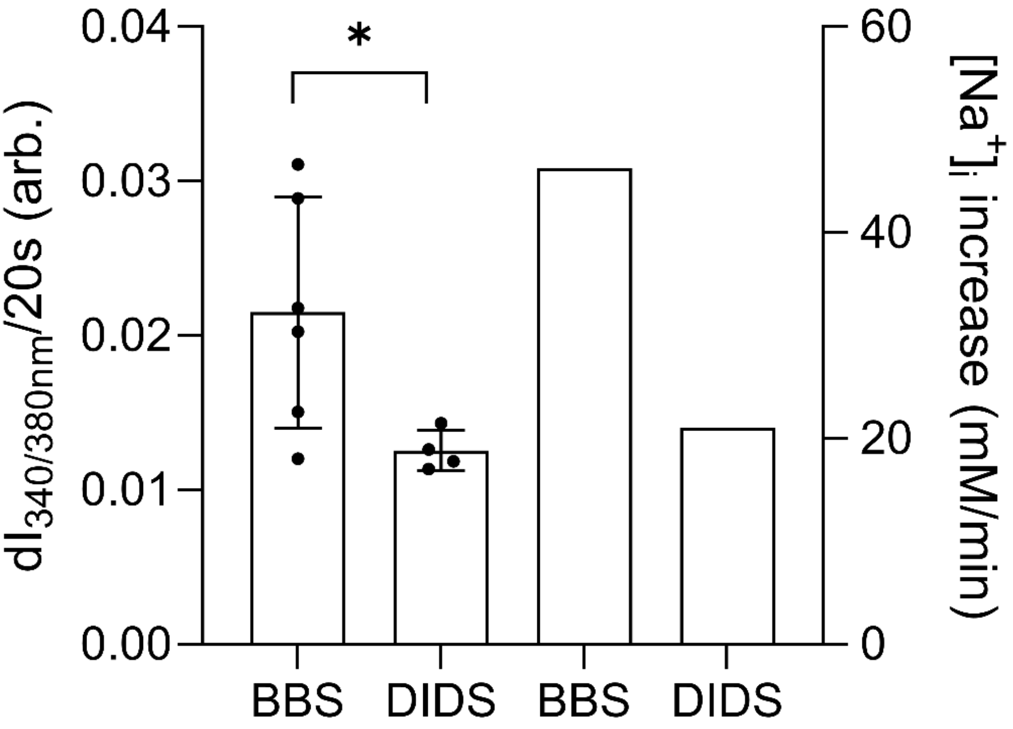


Supplemental Figure S1

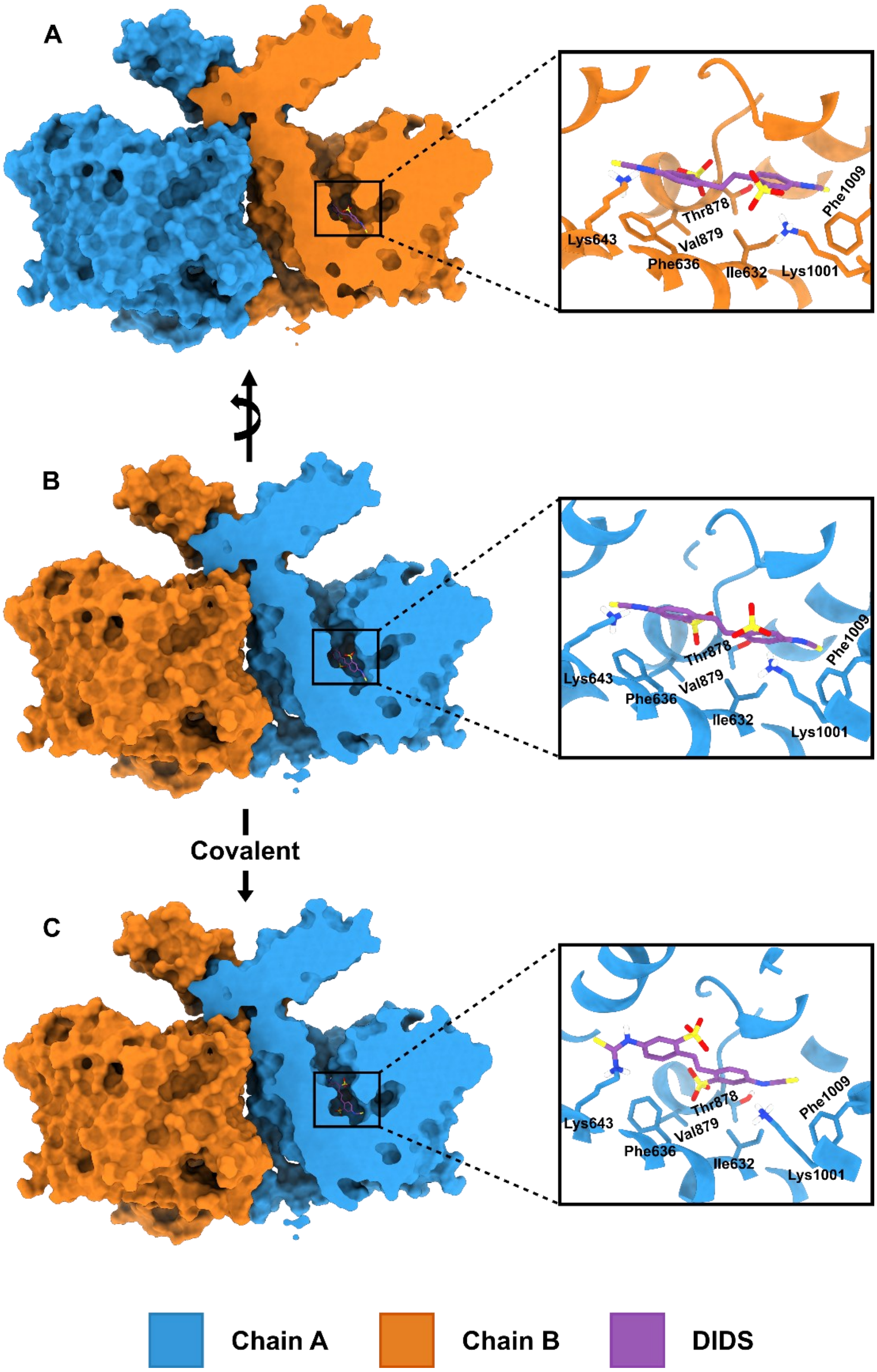


Supplemental Figure S2

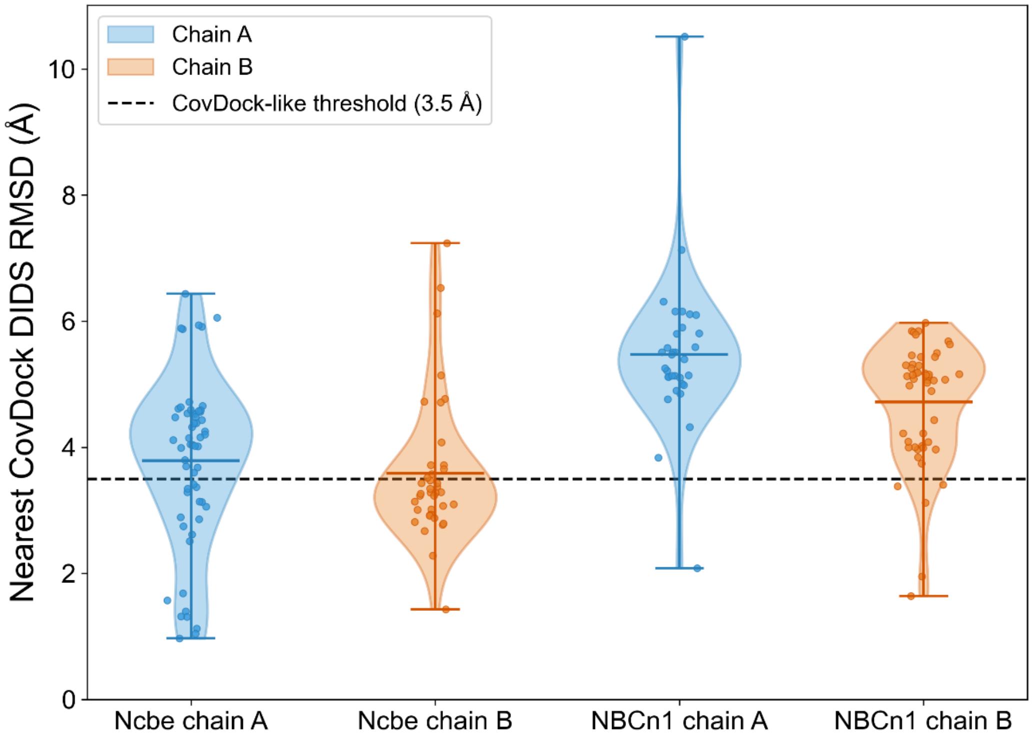


Supplemental Figure S3

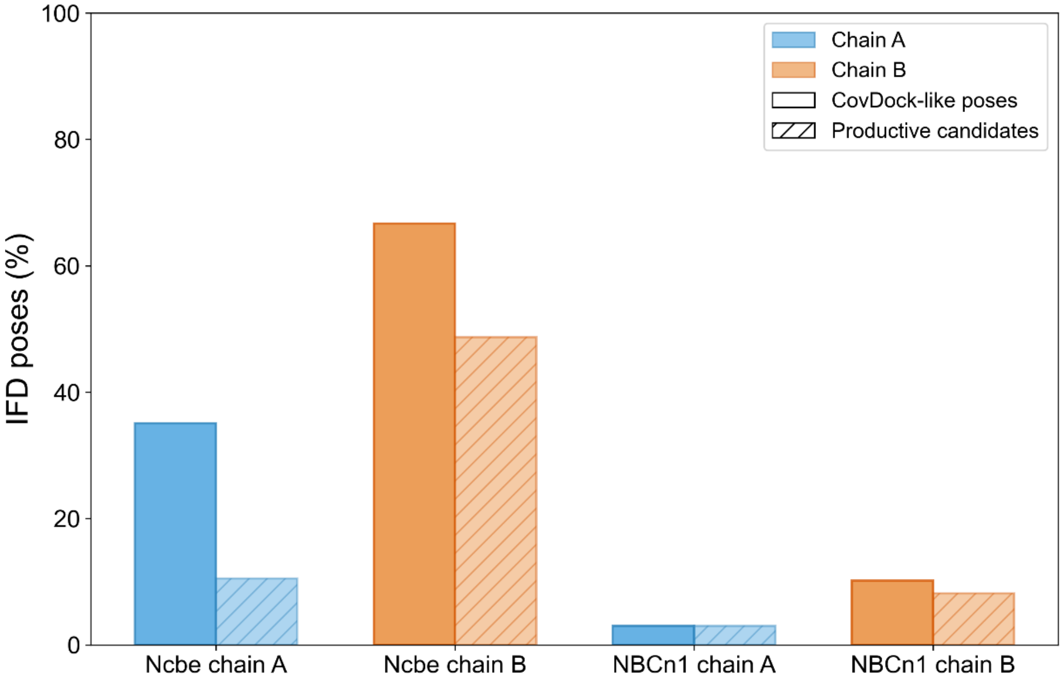


Supplemental Figure S4

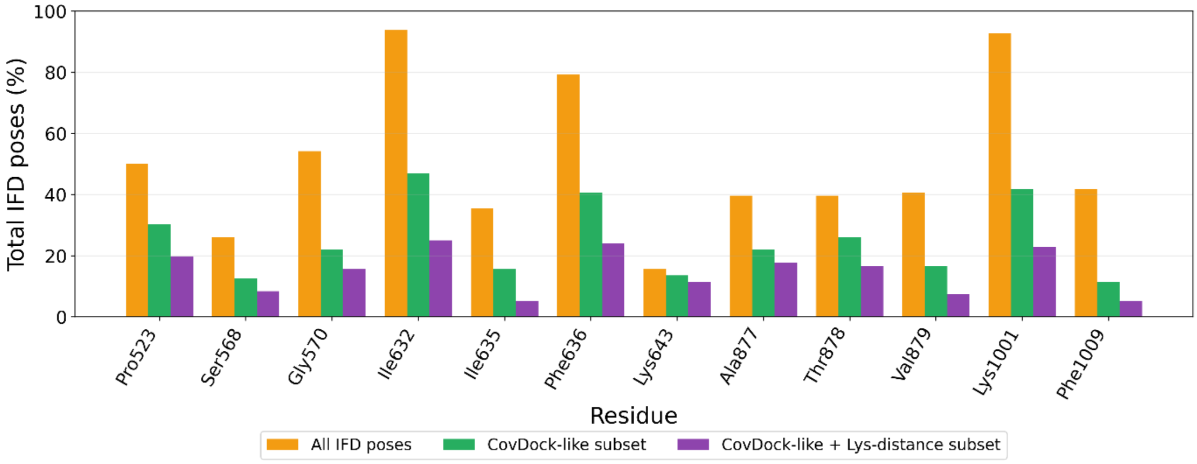


Supplemental Figure S5

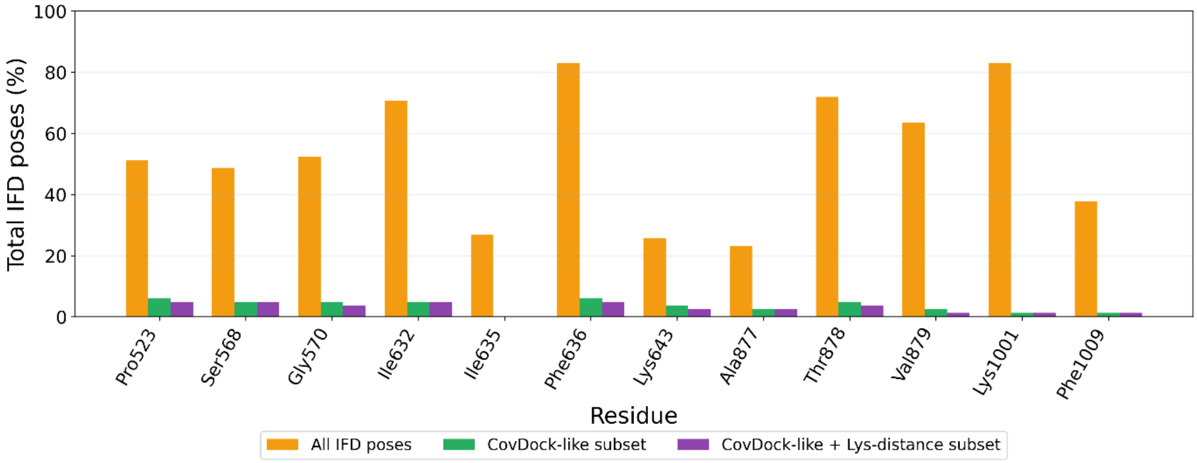


Supplemental Figure S6

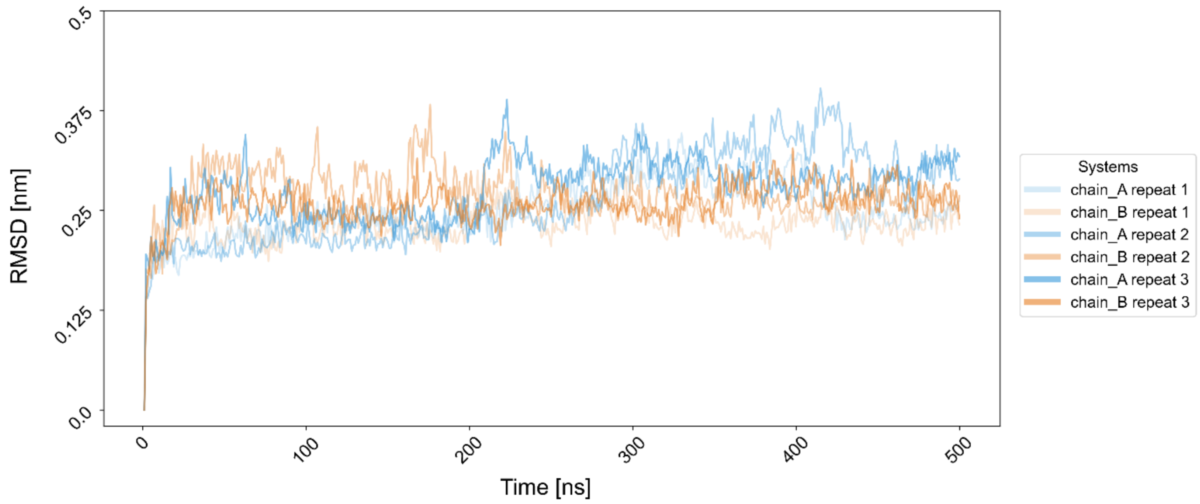


Supplemental Figure S7

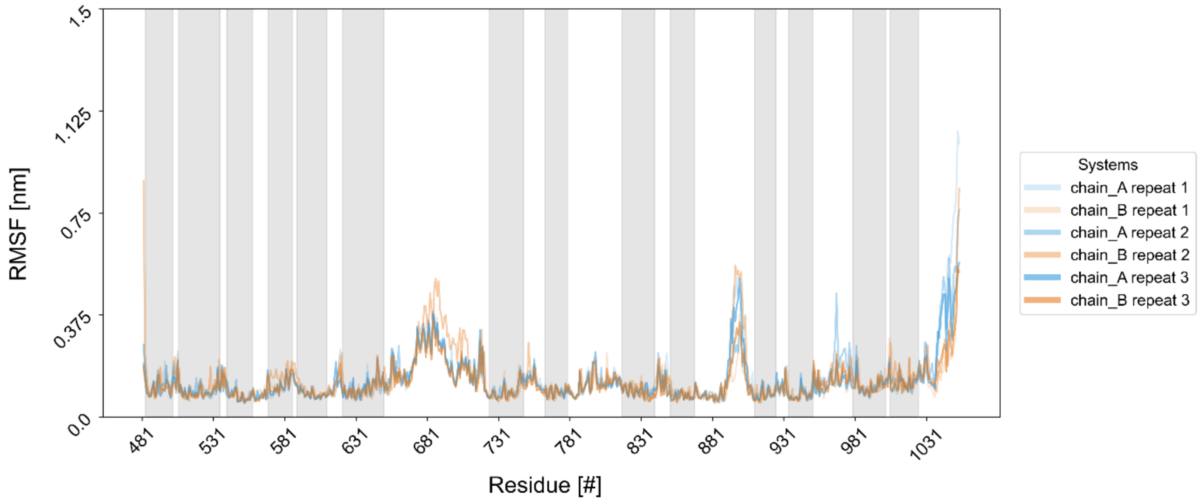


Supplemental Figure S8

Supplemental Table S1

| Protein | Chain | Cluster size | Docking score [kcal/mol] | IFDScore [kcal/mol] | MM-GBSA dG [kcal/mol] |
|---|---|---|---|---|---|
| Ncbe | A | 3 | -8.1 ± 0.7 | -32282.3 ± 14.3 | -59.2 ± 2.7 |
| Ncbe | A | 14 | -7.8 ± 0.8 | -32296.9 ± 16.4 | -50.5 ± 7.4 |
| Ncbe | A | 2 | -7.0 ± 0.7 | -32281.8 ± 30.3 | -48.0 ± 7.6 |
| Ncbe | A | 8 | -6.8 ± 0.9 | -32272.0 ± 16.7 | -43.9 ± 6.4 |
| Ncbe | A | 6 | -6.7 ± 0.5 | -32279.2 ± 7.8 | -46.4 ± 6.8 |
| Ncbe | A | 2 | -7.0 ± 0.1 | -32262.8 ± 9.1 | -47.1 ± 6.8 |
| Ncbe | A | 1 | -8.8 ± 0.0 | -32268.5 ± 0.0 | -49.2 ± 0.0 |
| Ncbe | A | 2 | -6.7 ± 0.0 | -32268.6 ± 6.4 | -45.1 ± 5.2 |
| Ncbe | A | 2 | -7.4 ± 0.1 | -32258.9 ± 18.5 | -46.5 ± 2.4 |
| Ncbe | A | 1 | -6.0 ± 0.0 | -32257.9 ± 0.0 | -47.9 ± 0.0 |
| Ncbe | A | 3 | -7.4 ± 0.1 | -32284.8 ± 3.4 | -43.3 ± 2.9 |
| Ncbe | A | 4 | -5.7 ± 0.9 | -32256.9 ± 14.9 | -35.5 ± 6.8 |
| Ncbe | A | 2 | -6.4 ± 0.3 | -32259.5 ± 12.4 | -40.9 ± 3.6 |
| Ncbe | A | 2 | -6.3 ± 0.3 | -32256.6 ± 18.3 | -38.0 ± 1.1 |
| Ncbe | A | 1 | -6.3 ± 0.0 | -32257.8 ± 0.0 | -38.8 ± 0.0 |
| Ncbe | A | 3 | -6.3 ± 0.4 | -32263.0 ± 5.3 | -34.0 ± 2.3 |
| Ncbe | A | 1 | -4.7 ± 0.0 | -32250.1 ± 0.0 | -27.9 ± 0.0 |
| Ncbe | B | 5 | -7.3 ± 0.9 | -32335.9 ± 16.9 | -42.8 ± 11.8 |
| Ncbe | B | 1 | -9.7 ± 0.0 | -32356.1 ± 0.0 | -52.0 ± 0.0 |
| Ncbe | B | 18 | -7.2 ± 0.8 | -32332.7 ± 14.8 | -39.3 ± 8.0 |
| Ncbe | B | 1 | -7.6 ± 0.0 | -32338.9 ± 0.0 | -47.6 ± 0.0 |
| Ncbe | B | 1 | -6.6 ± 0.0 | -32334.3 ± 0.0 | -46.9 ± 0.0 |
| Ncbe | B | 1 | -7.8 ± 0.0 | -32334.3 ± 0.0 | -45.8 ± 0.0 |
| Ncbe | B | 1 | -8.0 ± 0.0 | -32340.7 ± 0.0 | -42.2 ± 0.0 |
| Ncbe | B | 2 | -6.9 ± 0.3 | -32334.1 ± 15.8 | -39.0 ± 3.8 |
| Ncbe | B | 2 | -5.6 ± 0.4 | -32315.9 ± 12.2 | -36.7 ± 6.9 |
| Ncbe | B | 1 | -8.1 ± 0.0 | -32322.7 ± 0.0 | -39.2 ± 0.0 |
| Ncbe | B | 4 | -6.9 ± 0.3 | -32333.3 ± 9.1 | -36.7 ± 1.6 |
| Ncbe | B | 2 | -6.5 ± 0.2 | -32330.8 ± 5.2 | -30.9 ± 10.6 |

Supplemental Table S2

| Protein | Chain | Cluster size | Docking score [kcal/mol] | IFDScore [kcal/mol] | MM-GBSA dG [kcal/mol] |
|---|---|---|---|---|---|
| NBCn1 | A | 1 | -8.9 ± 0.0 | -40970.8 ± 0.0 | -62.6 ± 0.0 |
| NBCn1 | A | 2 | -8.1 ± 0.1 | -40962.3 ± 7.0 | -61.5 ± 1.5 |
| NBCn1 | A | 3 | -8.6 ± 0.4 | -40983.0 ± 13.9 | -56.7 ± 5.2 |
| NBCn1 | A | 1 | -8.4 ± 0.0 | -40961.3 ± 0.0 | -60.8 ± 0.0 |
| NBCn1 | A | 4 | -7.6 ± 0.6 | -40964.6 ± 19.7 | -57.1 ± 3.6 |
| NBCn1 | A | 1 | -9.0 ± 0.0 | -40963.7 ± 0.0 | -60.0 ± 0.0 |
| NBCn1 | A | 7 | -7.7 ± 0.6 | -40965.4 ± 22.4 | -54.3 ± 3.7 |
| NBCn1 | A | 3 | -7.7 ± 0.2 | -40934.5 ± 3.3 | -51.9 ± 0.9 |
| NBCn1 | A | 2 | -7.3 ± 0.6 | -40933.0 ± 19.8 | -46.2 ± 6.2 |
| NBCn1 | A | 2 | -6.7 ± 0.7 | -40935.7 ± 5.6 | -44.7 ± 2.8 |
| NBCn1 | A | 1 | -5.4 ± 0.0 | -40928.6 ± 0.0 | -45.8 ± 0.0 |
| NBCn1 | A | 1 | -5.4 ± 0.0 | -40919.8 ± 0.0 | -42.9 ± 0.0 |
| NBCn1 | A | 2 | -5.8 ± 0.5 | -40930.2 ± 8.0 | -40.4 ± 2.3 |
| NBCn1 | A | 1 | -6.4 ± 0.0 | -40920.6 ± 0.0 | -37.8 ± 0.0 |
| NBCn1 | A | 1 | -6.7 ± 0.0 | -40930.0 ± 0.0 | -36.9 ± 0.0 |
| NBCn1 | A | 1 | -4.8 ± 0.0 | -40922.4 ± 0.0 | -32.1 ± 0.0 |
| NBCn1 | B | 1 | -9.3 ± 0.0 | -40953.1 ± 0.0 | -64.0 ± 0.0 |
| NBCn1 | B | 5 | -8.6 ± 0.3 | -40926.0 ± 8.3 | -56.7 ± 4.0 |
| NBCn1 | B | 2 | -8.7 ± 0.6 | -40941.4 ± 5.2 | -58.5 ± 4.0 |
| NBCn1 | B | 5 | -7.6 ± 0.7 | -40925.6 ± 24.5 | -49.6 ± 8.0 |
| NBCn1 | B | 9 | -8.2 ± 0.5 | -40946.8 ± 11.9 | -56.2 ± 6.5 |
| NBCn1 | B | 2 | -9.0 ± 0.2 | -40935.9 ± 6.4 | -58.2 ± 0.9 |
| NBCn1 | B | 1 | -8.9 ± 0.0 | -40939.2 ± 0.0 | -57.9 ± 0.0 |
| NBCn1 | B | 1 | -8.6 ± 0.0 | -40928.1 ± 0.0 | -55.0 ± 0.0 |
| NBCn1 | B | 7 | -6.2 ± 0.4 | -40912.5 ± 5.4 | -39.0 ± 10.8 |
| NBCn1 | B | 2 | -6.4 ± 0.0 | -40923.3 ± 1.2 | -48.0 ± 0.3 |
| NBCn1 | B | 2 | -7.2 ± 0.5 | -40927.0 ± 2.8 | -46.6 ± 0.8 |
| NBCn1 | B | 1 | -6.4 ± 0.0 | -40894.3 ± 0.0 | -45.6 ± 0.0 |
| NBCn1 | B | 1 | -5.6 ± 0.0 | -40925.0 ± 0.0 | -45.2 ± 0.0 |
| NBCn1 | B | 3 | -5.5 ± 0.5 | -40905.9 ± 3.6 | -38.6 ± 6.2 |
| NBCn1 | B | 3 | -6.8 ± 0.1 | -40917.4 ± 6.7 | -42.5 ± 2.2 |
| NBCn1 | B | 1 | -5.8 ± 0.0 | -40887.2 ± 0.0 | -40.2 ± 0.0 |
| NBCn1 | B | 3 | -6.0 ± 0.4 | -40909.3 ± 2.1 | -29.5 ± 6.5 |

Supplemental Table S3

| Protein | Chain | Pose | CovDock affinity score | Combined docking score | Pre-reaction docking score | Post-reaction docking score | MM-GBSA dG [kcal/mol] |
|---|---|---|---|---|---|---|---|
| Ncbe | A | 1 | -4.8 | -4.8 | -5.1 | -4.5 | -17.4 |
| Ncbe | A | 2 | -4.0 | -4.0 | -4.9 | -3.2 | -23.7 |
| Ncbe | B | 1 | -2.3 | -2.3 | -4.3 | -0.3 | -8.6 |
| Ncbe | B | 2 | -3.8 | -3.8 | -4.3 | -3.4 | -25.7 |

Supplemental Table S4

| Protein | Chain | Pose | CovDock affinity score | Combined docking score | Pre-reaction docking score | Post-reaction docking score | MM-GBSA dG [kcal/mol] |
|---|---|---|---|---|---|---|---|
| NBCn1 | A | 1 | -1.9 | -1.9 | -4.5 | 0.6 | -18.7 |
| NBCn1 | A | 2 | -4.3 | -4.3 | -4.7 | -4.0 | -21.9 |
| NBCn1 | B | 1 | -4.0 | -4.0 | -4.5 | -3.4 | -22.6 |
| NBCn1 | B | 2 | -4.1 | -4.1 | -4.6 | -3.6 | -23.1 |
